# Chemotherapy synergizes with cancer vaccines and expands stem-like TCF1+CD8+ T cells

**DOI:** 10.1101/2023.12.18.572068

**Authors:** Laurine Noblecourt, Amanda Wicki, Vinnycius Pereira-Almeida, James McAuliffe, Emily Steffke, Silvia Panetti, Ramiro A. Ramirez-Valdez, Vineethkrishna Chandrasekar, Ashwin Jainarayanan, Adrian V.S. Hill, Benoit J. Van den Eynde, Carol Sze Ki Leung

## Abstract

Therapeutic cancer vaccines, whether based on neoantigens or shared antigens, will likely be given in the clinic together with the standard of care, which often comprises immune checkpoint blockade therapy and chemotherapy. It remains unclear, however, whether vaccines effectively synergize with chemotherapy. Here, we tested the combination of a heterologous prime-boost viral vector vaccine with chemotherapy (CarboTaxol) and anti-PD- 1. We show that this triple combination improves tumor control and survival in different murine tumor models. CarboTaxol, and also cyclophosphamide, acted as an immune adjuvant for the vaccines, enhancing tumor-specific CD8+ T-cell responses, irrespective of the presence of a tumor. These chemotherapies expanded stem-like T cell factor 1 (TCF1)+CD8+ T cells. Inhibition of the transcriptional activity of TCF1/β-catenin with a small molecule inhibitor abolished the immune adjuvant effect of CarboTaxol. This study sheds light on the new immunomodulatory roles of chemotherapies and holds promises for clinical testing of this combination strategy.

**Highlights:**

- The combination of CarboTaxol with viral vector cancer vaccines and anti-PD-1 promotes better tumor control, tumor clearance, and survival
- CarboTaxol increases TCF1 expression in CD8+ T cells and expands stem-like TCF1+CD8+ T cells
- CarboTaxol acts as an adjuvant for cancer vaccines irrespective of the presence of a tumor and this effect is mediated by TCF1/β-catenin activity

## Introduction

Therapeutic cancer vaccines have the potential to induce specific anti-tumor immune responses to reject tumors. However, their efficacy to date remains a challenge influenced by numerous factors ^1^. One major challenge is the immunosuppression induced by the tumor microenvironment (TME) ^2^. Combining cancer vaccines with immune checkpoint blockade (ICB) therapies targeting negative immune regulators such as Programmed Cell Death 1 (PD-1), or Cytotoxic T Lymphocyte-associated Antigen 4 (CTLA-4) can partly overcome the immunosuppression, leading to enhanced anti-tumor response in mice ^3^. While this combination therapy is being tested in the clinic ^4^, the combination of ICB and chemotherapy has become the first-line standard of care (SoC) in some types of cancer, such as non-small cell lung cancer (NSCLC). Chemotherapy drugs are usually cytotoxic, designed to kill rapidly dividing cancer cells, but some of them possess immune-activating properties that aid the immune system in eradicating tumor cells ^5^. Indeed, it has been reported that chemotherapy treatment, using carboplatin and paclitaxel (CarboTaxol), is efficient in depleting myeloid suppressor cells, leading to better efficacy of a long peptide vaccine ^6,7^. Moreover, personalized cancer vaccine with chemotherapy and anti-PD-1 has been tested in the clinic as first-line therapy for advanced NSCLC, indicating it is safe and practical ^8,9^, but the anti-tumor efficacy of this triple combination therapy remains to be determined.

Key to the efficacy of anti-tumor immunotherapies are stem-like CD8+ T cells, which play an important role in sustaining durable T cell responses ^10,11^. These cells are characterized by their self-renewal, differentiation and multipotent capacity, often express high levels of T Cell Factor 1 (TCF1, encoded by *Tcf7*), which is a key transcription factor of the canonical Wingless/Integration 1 (WNT)/β-catenin signaling pathway. TCF1 has a pivotal role in T-cell development, stemness and memory formation ^12^. Ablation of *Tcf7* impairs secondary expansion of virus-specific CD8+ T cells upon LCMV reinfection, highlighting the role of TCF1 in the generation of long-lived CD8+ memory T cells ^13^. Moreover, ectopic TCF1 expression promotes the formation of TCF1+ stem-like precursor T cells in a mouse model of melanoma, leading to tumor control ^14^. However, the impact of various cancer therapies on TCF1 expression are still not well understood.

Here, we investigated the combination of viral vector vaccines, chimpanzee adenovirus (ChAdOx1) and Modified Vaccinia Ankara (MVA), with CarboTaxol and anti-PD-1 using a tumor model expressing P1A, the mouse counterpart of Melanoma Antigen Gene (MAGE)-type antigens. We show that this triple combination therapy enhances immunogenicity and has superior anti-tumor efficacy. CarboTaxol improves vaccine-induced P1A-specific CD8+ T cell response and expands TCF1+CD8+ T cells in a tumor-independent manner. Furthermore, inhibiting the transcriptional activity of TCF1/β-catenin abrogates the adjuvant effect of CarboTaxol, underscoring the pivotal role of TCF1 in priming and expanding CD8+ T cells against tumor antigens.

## Results

### Combination of ChAdOx1/MVA vaccines, anti-PD-1 and CarboTaxol improves tumor control, survival, and immunogenicity

We implanted DBA/2 mice with 15V4T3 tumors, which naturally express the P1A antigen. To assess the efficacy of combined ChAdOx1/MVA, anti-PD-1 and CarboTaxol treatment, hereafter referred to as the triple combination therapy, we vaccinated the mice, when tumors were palpable, with P1A-encoding ChAdOx1, and treated them with CarboTaxol at the same time. We boosted the mice with P1A-encoding MVA a week later. Control mice were sham-vaccinated with ChAdOx1 or MVA encoding an irrelevant protein. The anti-PD-1 treatment was given every third day for a total of 3 injections after the boost (Figure 1A).

**Figure 1:**
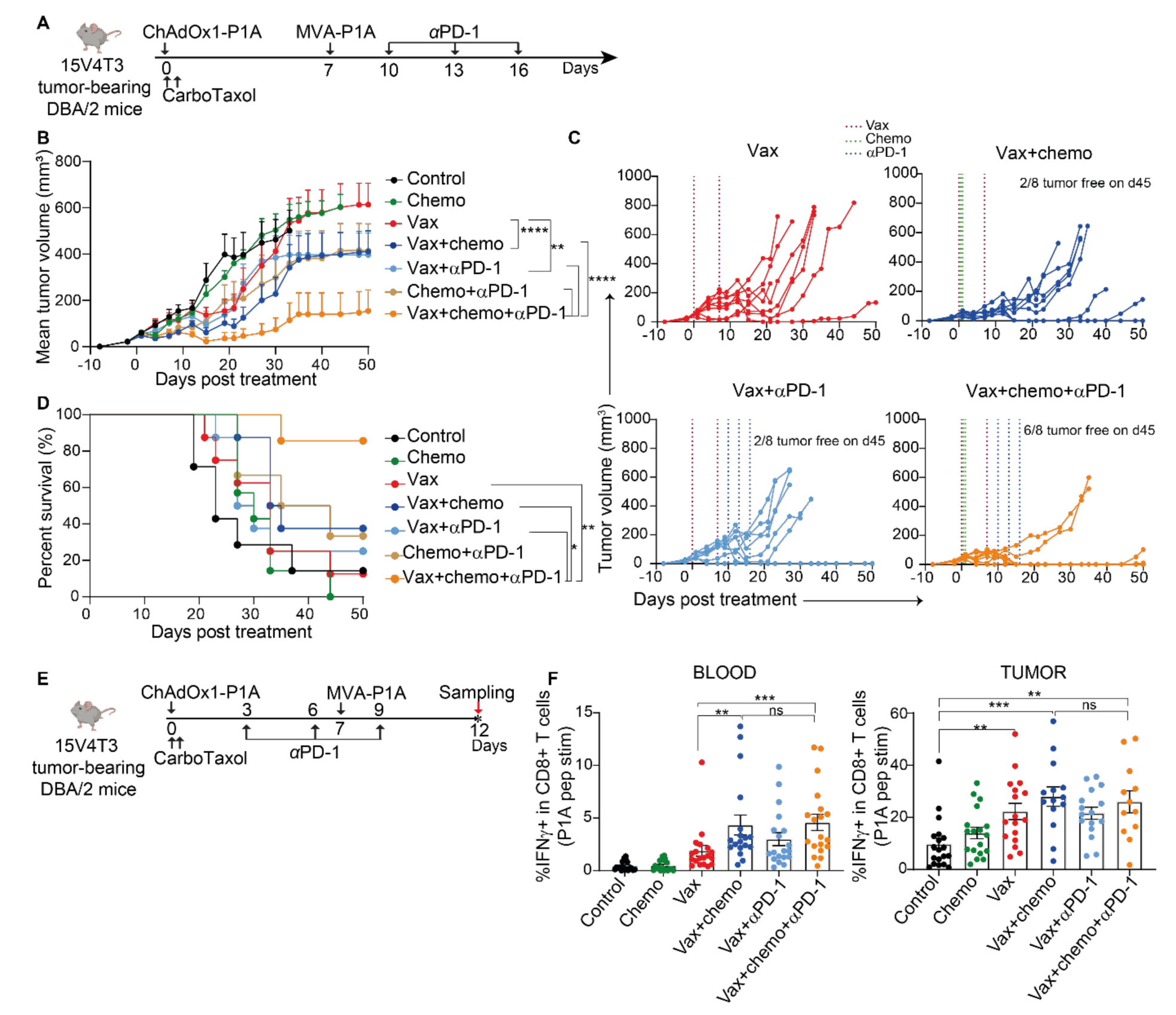
Triple combination of ChAdOx1/MVA-P1A vaccines, anti-PD-1 and CarboTaxol improves tumor control, survival and immunogenicity. **(A)** Schematic representing timeline of experiment. DBA/2 mice were implanted with 15V4T3 cells to initiate tumor growth. Eight to 11 days after tumor implantation, mice were vaccinated with ChAdOx1-P1A and treated with carboplatin and paclitaxel (CarboTaxol). The mice were then vaccinated with MVA-P1A and treated with anti-PD-1 (αPD-1 in scheme and graphs) every third day for 3 injections. Mice were vaccinated with ChAdOx1/MVA-DPY in the control group. **(B-D)** Mean (+ SEM) tumor growth **(B)**, individual tumor growth curves **(C)** and survival **(D)** are shown. Data are representative of at least 3 experiments (n= 6 to 8 mice per group). Statistically significant differences in tumor volume between groups were determined by a two-way ANOVA followed by Tukey’s post hoc test and statistical differences in survival were determined by a Log Rank test. **(E)** An alternative treatment scheme for sample harvest was used where the anti-PD-1 was administered 3 days after ChAdOx1 vaccination. On day 12 post treatment, PBMCs and tumor cells were stimulated with P1A peptides and the proportion of IFNγ-producing P1A-specific CD8+ cells was determined by flow cytometry. **(F)** Average percentage of IFNγ+ producing CD8+ T cells in blood (n = 18 to 19 mice) and tumor (n= 10 to 19 mice) are shown as mean ± SEM. Data are representative of at least 3 independent experiments with each symbol representing an individual mouse. Statistically significant differences between vaccinated groups were determined by a Mann-Whitney test. ns non-significant, * p ≤ 0.05, ** p ≤ 0.01, *** p ≤ 0.001 **** p ≤ 0.0001.

We observed a significant improvement in tumor growth control in mice receiving the triple combination therapy compared to the vaccines alone, vaccines + CarboTaxol or vaccines + anti-PD-1 (Figure 1B). Interestingly, we also observed significantly improved tumor growth control in the vaccines + CarboTaxol group, compared to the vaccines alone treatment group (Figure 1B). Remarkably, with the triple combination therapy, six of eight mice were tumor-free on day 45 after treatment (Figure 1C). The enhanced efficacy of the triple combination was also reflected in mouse survival. At day 50 post treatment, 85% of the mice in the triple combination group were still alive, whilst less than 20% of the mice survived in the vaccines alone group (Figure 1D). Altogether, these data show that the triple combination has superior tumor growth control and survival compared to the other treatment regimes.

We also investigated this triple combination therapy in HPV16 E6 and E7 positive TC- 1 mouse tumor model. To this end, C57BL/6 mice were implanted with TC-1 cells expressing the E6 and the E7 tumor antigens. Mice were vaccinated with ChAdOx1/MVA-E6/E7 and treated with anti-PD-1 and CarboTaxol as represented in the scheme (Supplemental Figure 1A). Adding just CarboTaxol to the vaccines already improved tumor control over the vaccines alone or chemo alone in this model, indicating an essential role of CarboTaxol in improving the anti-tumor effect of the vaccines (Supplemental Figure 1 B-C). Nevertheless, individual tumor growth data highlighted that the triple combination of ChAdOx1/MVA-E6/E7, CarboTaxol, and anti-PD-1 tended to induce superior tumor regression (Supplemental Figure 1D). In line with the improved tumor regression, the triple combination group displayed the best survival and was significantly better than the vaccines alone group (Supplemental Figure 1C).

Chemotherapy can have immunomodulatory effects and can be combined with immunotherapy strategies to enhance their efficacy ^5^. To understand the impact of CarboTaxol on vaccine-induced P1A-specific anti-tumor T cells, we treated mice according to the scheme shown in Figure 1E. We administered anti-PD-1 three days after the prime in order to include this treatment before the harvest of tumor and blood samples on day 12 post treatment, when tumor regressed dramatically in responders. We stimulated peripheral blood mononuclear cells (PBMCs) or tumor-derived cell suspensions with overlapping P1A peptides before evaluating the fraction of cytokine-producing P1A-specific CD8+ T cells by flow cytometry. The addition of CarboTaxol to vaccination significantly increased the proportion of IFNγ-producing P1A-specific CD8+ T cells in the blood compared to the vaccines alone group (Figure 1F). The vaccines + CarboTaxol and the triple combination groups displayed the highest tumor infiltration of P1A-specific CD8+ T cells, though it did not reach significance compared to the vaccines alone. It is worth mentioning that these groups also presented high responders with tumor regression which did not allow for the analysis of all tumor samples. Importantly, the addition of anti-PD-1 did not further increase the magnitude of the P1A-specific response in the blood or in the tumors, as the response of the vaccines + CarboTaxol and the triple combination groups were comparable (Figure 1F). However, the addition of anti-PD-1 seemed to support better P1A-specific CD8+ responses in the blood at later time points (Supplemental Figure 2, A-B). Importantly, the P1A-specific CD8+ T cell response in the blood was positively correlated with tumor control, highlighting the importance of antigen-specific responses (Supplemental Figure 2C).

### CarboTaxol enables better flexibility in the administration window of anti-PD-1 as part of the triple combination strategy

To avoid delay in the standard of care (SoC) treatment, it is advisable for patients to start receiving the SoC treatment while undergoing testing for expression of the cancer antigen targeted with the vaccines, or during preparation of the personalized vaccine. Therefore, anti-PD-1 will likely be given prior to vaccination in clinical testing as it is part of the SoC. Hence, we tested a new administration strategy in our model with anti-PD-1 given prior to the vaccines (Vax+pre-vax αPD-1). This new schedule was compared with the previously evaluated post-prime schedule of anti-PD-1 treatment, starting 3 days post-prime for 3 injections every third day (Vax+post-vax αPD-1) (Figure 2A). While administering anti-PD-1 after vaccination improved tumor control over the vaccines alone, the new schedule of pre-vax anti-PD-1 did not (Figure 2B). This indicates a loss of synergy between the vaccines and anti-PD-1 when the latter is administered prior to vaccination, in line with previous findings demonstrating that PD-1 blockade prior to vaccination decreased the beneficial effects of a peptide vaccine on tumor control ^15^. Interestingly however, both triple combination groups showed a similar tumor control (Figure 2B) and survival (Figure 2C), which were significantly better than the vaccines + anti-PD-1 and vaccines alone groups. These data suggested that CarboTaxol rescued the loss of synergy observed in the vaccines + pre-vax anti-PD-1 combination, promoting better tumor control (Figure 2B) and survival (Figure 2C) than vaccines alone or vaccines + anti-PD-1 groups.

**Figure 2:**
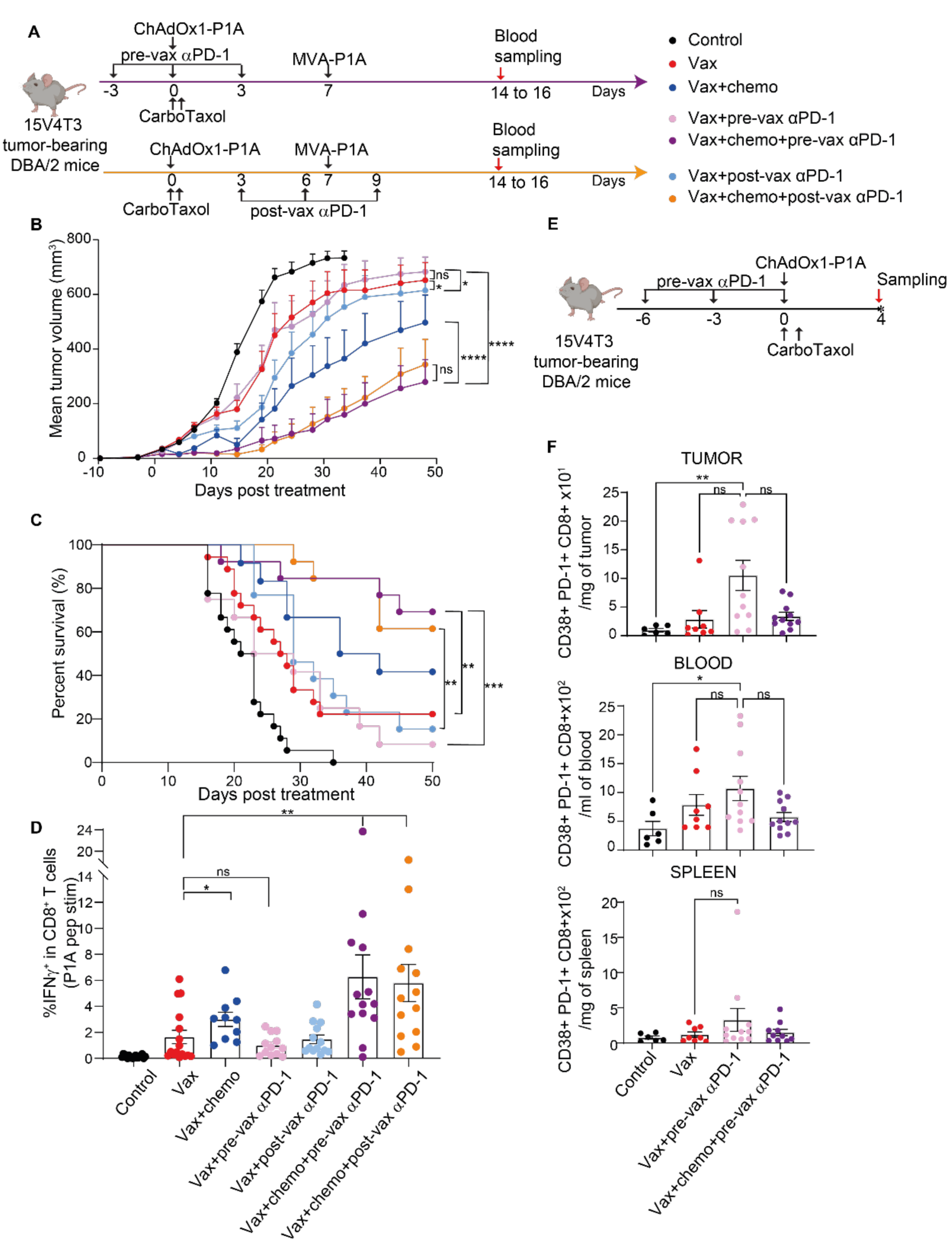
CarboTaxol enables better flexibility in the administration window of anti-PD-1 as part of the triple combination strategy. **(A)** Schematic representing timeline of experiment. DBA/2 mice were implanted with 15V4T3 and treated with the pre-vax anti-PD-1 regimen starting prior to vaccination with ChAdOx1-P1A. Mice were also treated with CarboTaxol and, 7 days later, were vaccinated with MVA-P1A. Some mice were treated with anti-PD-1 starting on day 3 post vaccination, referred as post-vax anti-PD-1. **(B)** Mean tumor growth is shown. Data are presented as mean tumor volume (mm^3^) + SEM and statistically significant differences between groups were determined by two-way ANOVA followed by Tukey’s post hoc tests. **(C)** Percentage of mouse survival in each group is shown and statistical differences in survival were determined by a Log Rank test. For (B) and (C) data were pooled from 2 to 3 independent experiments (n= 10 to 18 mice per groups). **(D)** The percentage of IFNγ producing cells in CD8+ T cells in the blood are shown as mean ± SEM. Data were pooled from 2 to 3 independent experiments (n= 10 to 18 mice per group). Significant differences between the relevant groups were determined by Mann-Whitney test. **(E)** Schematic representing timeline of experiment. Mice were treated with the pre-vax anti-PD-1 regimen vaccinated with ChAdOx1-P1A or PBS sham and administered CarboTaxol. **(F)** Mean cell numbers ± SEM of CD38+PD-1+ CD8+T cells are shown in tumor, blood and spleen. Data were pooled from 2 independent experiments (n= 6 to 11 mice per group) and significant differences were determined by a Kruskal-Wallis test with Dunn’s multiple comparisons test. ns non-significant, * p ≤ 0.05, ** p ≤ 0.01, *** p ≤ 0.001, **** p ≤ 0.0001.

We then evaluated the effects of anti-PD-1 timing on the magnitude of the P1A- specific CD8+ T cell response in the blood. The magnitude of IFNγ-producing CD8+ T cells, after P1A peptide stimulation, was similar between mice receiving the vaccines alone or in combination with pre-vax anti-PD-1, while the vaccines + CarboTaxol and both triple combinations groups significantly increased the magnitude of P1A-specific CD8+ T cells compared to the vaccines alone (Figure 2D). This is in line with the tumor growth and survival data and highlights the importance of CarboTaxol in rescuing the loss of synergy when combining vaccines + pre-vax anti-PD-1.

It has been demonstrated that anti-PD-1 prior to vaccination can induce dysfunctional sub-primed CD38+PD-1+CD8+ T cells, which are more apoptotic and less prone to produce IFNγ ^15^. Therefore, we analyzed the level of CD38+PD-1+CD8+ T cells, at 4 days post treatment in multiple tissues (Figure 2E). Consistent with these findings, we observed a trend towards an expansion of CD38+PD-1+ cells in the tumor, spleen and blood following the vaccine+pre-vax anti-PD-1 regimen compared to the vaccine alone group, although it did not reach statistical significance (Figure 2F). Interestingly, when CarboTaxol was combined with pre-vax anti-PD-1 and ChAdOx1, CD38+PD-1+CD8+ T cells were not expanded, highlighting a potential effect of CarboTaxol in promoting T-cell fitness (Figure 2F).

### CarboTaxol depletes CD11b+Ly6G+ cells

Previous studies have shown that CarboTaxol can deplete CD11b+Gr1 high (^hi^) cells, and therapeutic vaccination during this myeloid depletion promoted better tumor control and survival ^6,7^. Hence, we questioned whether the depletion of potential suppressive myeloid cells, also known as myeloid-derived suppressor cells (MDSCs) or pathologically activated neutrophils, was occurring in our model, contributing to the observed synergy between vaccination and CarboTaxol. 15V4T3 tumor-bearing mice were treated with CarboTaxol and vaccinated as described in Figure 3A. Myeloid populations were investigated in different organs 3-4 days post treatment. Two main myeloid populations consisting of CD11b+Gr1^hi^ cells and CD11b+Gr1^lo^ cells were characterized. CD11b+Gr1^hi^ cells, which consist of 99% Ly6G+ cells (Figure 3B) can be characterized as neutrophils or polymorphonuclear (PMN)- MDSCs depending on their suppressive functions ^16^. The CD11b+Gr1^lo^ population is more heterogeneous and consists of Ly6C negative (^neg^), intermediate (^int^) and high (^hi^) populations (Figure 3B). CD11b+Gr1^lo^Ly6C^hi^ cells can be characterized as monocytic (M)-MDSCs if they display suppressive functions ^16^. CarboTaxol alone or in combination with ChAdOx1 significantly depleted CD11b+Gr1^hi^Ly6G+ cells, and to a lesser extent, the CD11b+Gr1^lo^ population (Figure 3B), in blood, bone marrow (BM), spleen and tumor (Figure 3 C-D). The CD11b+Gr1^lo^Ly6C^hi^ cells, potential M-MDSCs, were only slightly decreased by CarboTaxol (Supplemental Figure 3 A-B). Moreover, the depletion of CD11b+Gr1^hi^Ly6G+ by CarboTaxol was also observed in the TC-1 tumor model in C57BL/6 mice (Supplemental Figure 3 C-D).

**Figure 3:**
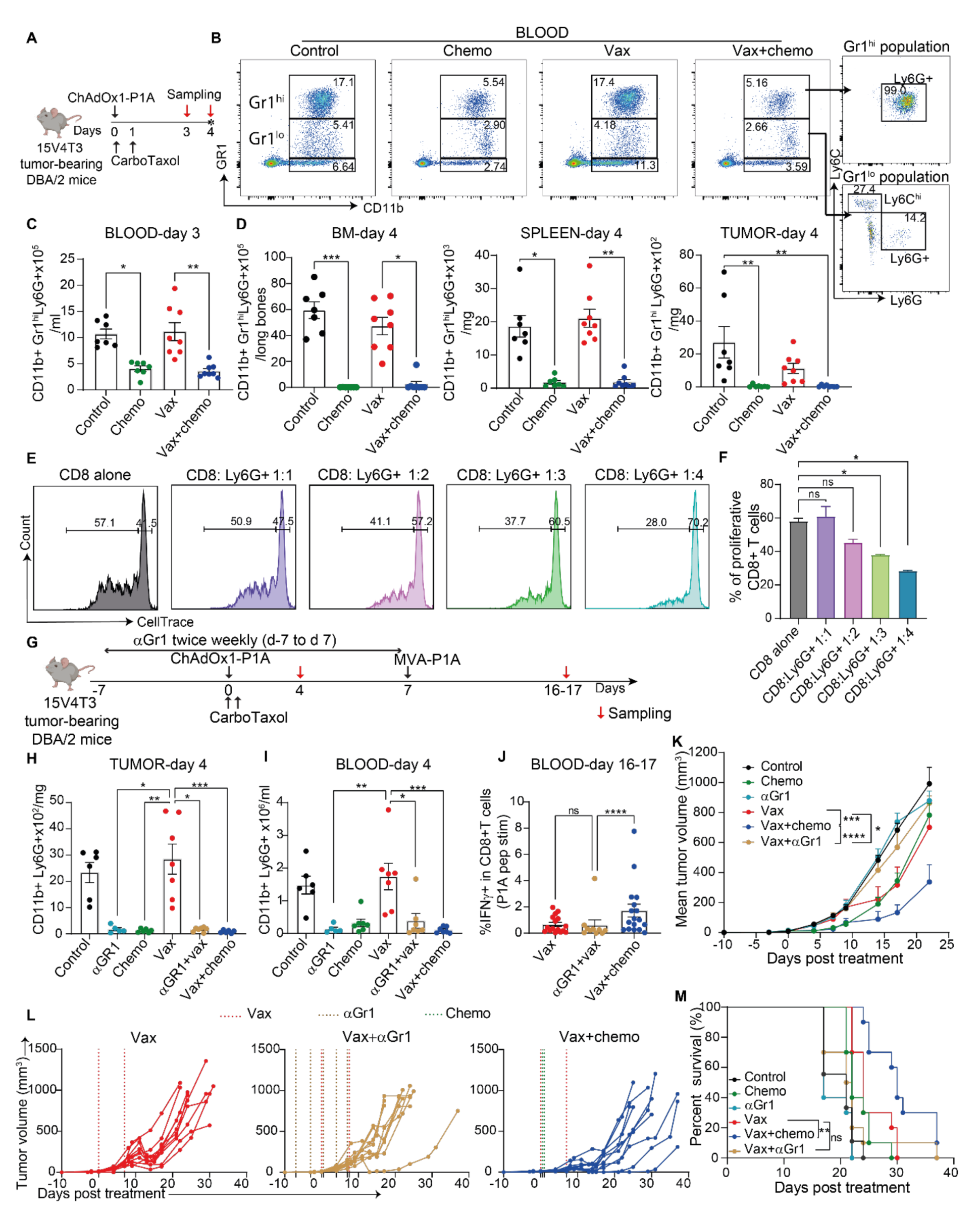
CarboTaxol depletes CD11b+Ly6G+ cells. **(A)** Schematic representing timeline of experiment. Eleven days after tumor implantation with 15V4T3, DBA/2 mice were vaccinated with ChAdOx1-P1A or PBS sham and treated with CarboTaxol. On day 3 post-prime, the blood was harvested, and 24 h later, spleen, BM and tumor were harvested. **(B)** CD11b+Gr1^hi^ and ^lo^ from the blood are shown, comparing the different treatment groups. The composition of these populations in Ly6G+ and Ly6C+ cells is indicated. (**C**-**D**) Numbers of CD11b+ Gr1^hi^ Ly6G+ are shown in the blood (**C**), the BM, the spleen and the tumor (**D**). The data are representative of 2 independent experiments and mean ± SEM for each group is shown. Each symbol represents an individual mouse (n=7 to 8 mice per group). **(E)** CD11b+Ly6G+ cells from spleen of tumor bearing mice and CD8+ T cells were co-cultured at different ratio and proliferation of CD8+ T cells upon stimulation with CD3 and CD28 antibodies was assessed 72 h later. Gating of proliferative and non-proliferative CD8+ T cells at 72h is shown. **(F)** Percentage of proliferative cells are shown as mean ± SEM. The data were pooled from 2 independent replicates. **(G)** DBA/2 mice implanted with 15V4T3 cells were vaccinated with ChAdOx1-P1A or PBS sham 10 to 14 days after tumor implantation. Additionally, mice were treated with either CarboTaxol or with anti-Gr1 antibody twice a week starting 7 days prior to vaccination. Mice were boosted with MVA-P1A. (**H-I**) The numbers of CD11b+Ly6G+ cells in the tumor (**H**) and blood (**I**) are shown as mean ± SEM. Data were pooled from 2 independent experiments (n= 6 to 7 mice). For (C), (D), (F), (H), (I) statistics were determined by a Kruskal-Wallis test with Dunn’s multiple comparisons test. **(J)** The proportion of IFNγ+ P1A-specific CD8+ T cells in the blood is shown as mean ± SEM. Data were pooled from at least 2 independent experiments (n= 9 to 17 mice) and statistically significant differences were determined by a Mann-Whitney test. **(K)**Mean tumor growth (mm^3^) + SEM is shown and is representative of 2 independent experiments (n=5 to 6 mice per group). Statistically significant differences were determined by two-way ANOVA followed by Tukey’s post hoc test. **(L)**Individual tumor growth curves are shown. (**M**) Percentage of survival is shown. For (L-M) data are pooled from 2 independent experiments (n= 9 to 10 mice). Statistical differences in survival data (M) were determined by a Log Rank test. ns non-significant, * p ≤ 0.05, ** p ≤ 0.01, *** p ≤ 0.001 **** p ≤ 0.0001.

To test whether the CD11b+Gr1^hi^Ly6G+ population displays immunosuppressive function in 15V4T3 tumor model, we investigated the impact of these cells on CD8+ T-cell proliferation. CD8+ T cells were isolated from the spleen of tumor-free mice and stimulated with CD3/CD28 antibodies in the presence of CD11b+Ly6G+ cells from spleen of 15V4T3 tumor-bearing mice at different ratios. We observed a 2-fold decrease in proliferating CD8+ T cells when Ly6G+ cells were added at a high ratio of 1:4 (Figure 3 E-F). These data suggest that the Ly6G+ population in 15V4T3 may comport PMN-MDSCs but with limited suppressive function. Moreover, a significant systemic expansion of suppressive myeloid populations is usually observable upon tumorigenesis in patients and animal models. However, the 15V4T3 tumor model did not display an expansion of CD11b+GR1^hi^Ly6G+ cells compared to tumor-free mice (Supplemental Figure 3E). Notably, the depletion of CD11b+Gr1^hi^Ly6G+ cells after CarboTaxol treatment was transient and was not observed on day 12 post treatment (Supplemental Figure 3 F-G). Therefore, to determine the importance of myeloid cell depletion in our model, and whether it could explain the increased vaccine efficacy with CarboTaxol, we depleted CD11b+Ly6G+ cells by injecting DBA/2 mice twice weekly with an anti-Gr1 antibody starting on day 3 post tumor implantation, and we combined it with vaccination (Figure 3G). Depletion of CD11b+Ly6G+ cells with anti-Gr1 in the blood and tumor was comparable to CarboTaxol, regardless of concomitant vaccination (Figure 3 H-I). However, mice treated with anti-Gr1 + vaccines did not show an increase in the magnitude of P1A- specific CD8+ T cells as observed in the vaccines + CarboTaxol group (Figure 3J). Furthermore, combining anti-Gr1 with the vaccines did not improve tumor control (Figure 3 K-L), nor survival (Figure 3M) compared to the vaccines alone group, whilst the vaccines + CarboTaxol treatment displayed a significantly better tumor control and survival. The same experiment was performed using anti-Ly6G antibody, which surprisingly appeared less efficient at depleting CD11b+Ly6G+ cells, and ultimately displayed no effect on tumor control nor vaccine immunogenicity (Supplemental Figure 3 H-K). Therefore, this data suggest that there is potentially another mechanism contributing to the synergy between ChAdOx1/MVA vaccination and CarboTaxol.

### The adjuvant effect of CarboTaxol is tumor-independent

Chemotherapeutic agents can be immunomodulatory through different mechanisms that can be separated into two major groups: tumor-independent mechanisms, which directly impact the immune system, or tumor-dependent processes, such as immunogenic cell death (ICD), which stimulate dendritic cells (DCs) and promote CD8+ T-cell activation through the release of danger signals following tumor cell death ^5^. To distinguish between these two mechanisms, we asked whether the increase in vaccine-induced P1A-specific CD8+ T cells was maintained in tumor-free mice. We vaccinated tumor-free DBA/2 mice, and administered CarboTaxol on the same day as the prime vaccine (same day chemo), as we did in tumor-bearing mice. Given the absence of tumor, the interval between the prime and boost was adjusted to 4 weeks apart to have an optimal immune response ^3^. This allowed us to also investigate different schedules in which CarboTaxol was given either 7 days prior to the prime (early chemo), 14 days after prime (late chemo) or together with the boost (chemo with boost) (Figure 4A). The magnitude of the P1A-specific CD8+ T-cell response was evaluated based on the proportion of IFNγ-producing CD8+ T cells post-prime and post-boost. Both were significantly increased with the vaccines + same day chemo treatment compared to vaccines alone, demonstrating a long-lasting tumor-independent adjuvant effect of CarboTaxol (Figure 4B, vax+ same day chemo). In contrast, when CarboTaxol was administered on a different day than the ChAdOx1 vaccine, the adjuvant effect was lost (Figure 4B, vax+early chemo, vax+late chemo). Interestingly, when CarboTaxol was administered together with the boost vaccination, it also increased the magnitude of the P1A-specific CD8+ T-cell response compared to the vaccines alone (Figure 4B, vax+chemo with boost). Upon subsequent challenge with a P1A-expressing tumor, mice treated with vaccines + same day chemo displayed better tumor protection and survival as compared to vaccines alone (Figure 4 C-D), confirming the importance of CarboTaxol in promoting a strong and long-lasting P1A-specific CD8 response upon vaccination. Remarkably, when given together with a different vaccine platform comprising of a self-assembling peptide nanoparticle TLR-7/8 agonist (SNP) ^17^, CarboTaxol also increased the magnitude of the P1A-specific response post-boost (Supplemental Figure 4 A-B), indicating the adjuvant effect of CarboTaxol is not restricted to viral-vector vaccine platforms. These results highlight the vaccine adjuvant effect of CarboTaxol in a tumor-independent manner, and therefore exclude a role of ICD in this effect.

**Figure 4:**
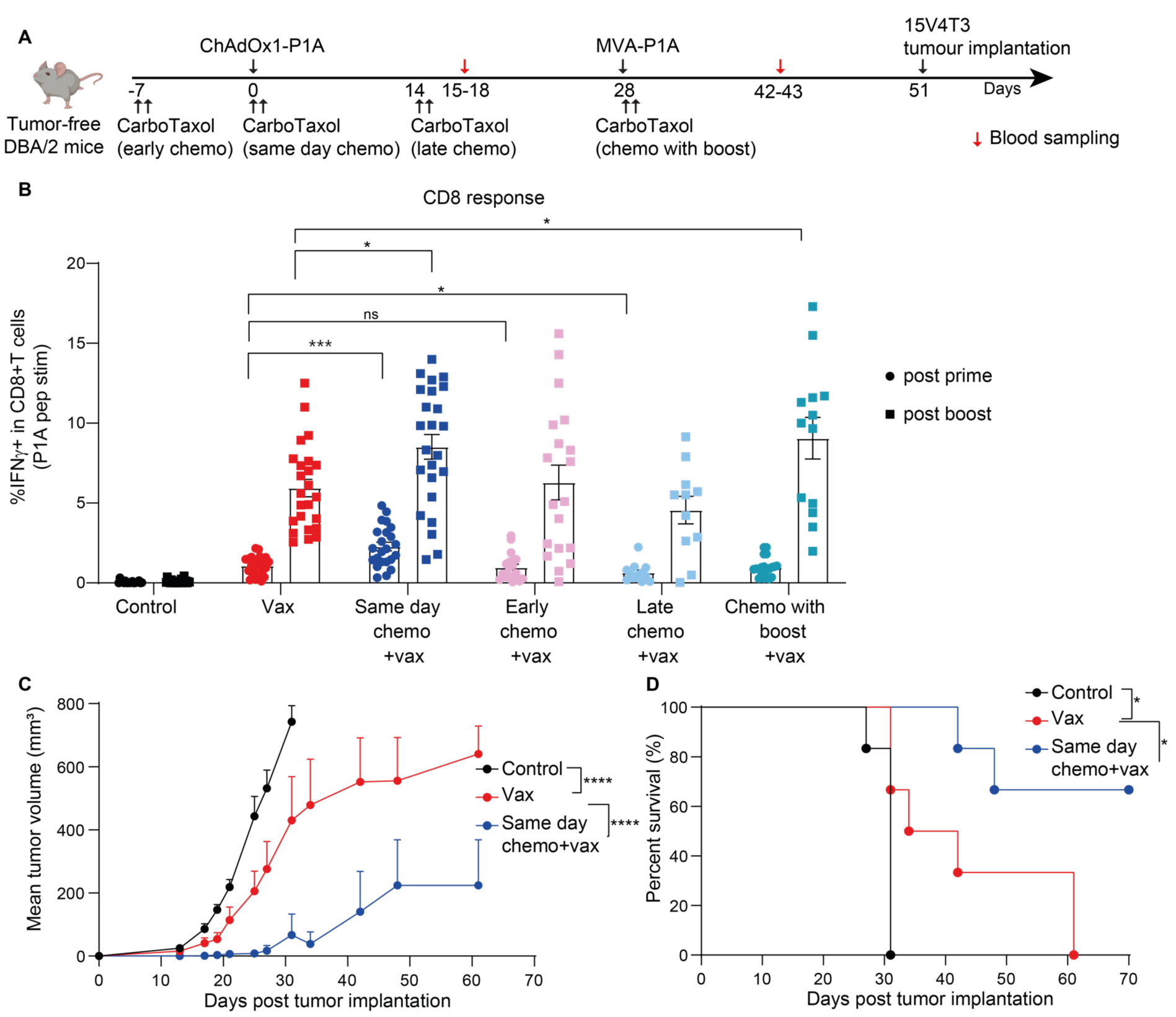
The adjuvant effect of CarboTaxol is tumor independent. **(A)** Schematic representing timeline of experiment. DBA/2 mice were vaccinated with ChAdOx1-P1A or PBS sham. Twenty-eight days later the mice received MVA-P1A. Mice were also treated with CarboTaxol given 7 days before the prime (early chemo), or together with the prime (same day chemo), or 14 days after the prime (late chemo), or together with the boost (chemo with boost). 15-18 days post-prime and 14-15 days post-boost, blood was harvested and the P1A-specific CD8 response was evaluated. **(B)** The proportion of IFNγ-producing P1A-specific CD8+ T cells are shown as mean ± SEM post-prime and post-boost. Data are pooled from at least 2 independent experiments with each symbol representing an individual mouse (n= 12 to 24 mice per group) and statistically significant differences between groups were determined by a Mann-Whitney test. (**C**-**D**) 3 to 6 weeks post-boost, mice were implanted with 15V4T3 cells and monitored to evaluate tumor progression. Mean tumor growth (mm^3^) + SEM (**C**) and percentage of survival (**D**) are shown (n= 6 mice per group). Statistically significant differences between groups for (C) were determined by two-way ANOVA followed by Tukey’s post hoc test and statistical differences in survival data (D) were determined by a Log Rank test. ns non-significant, * p ≤ 0.05, *** p ≤ 0.001.

We also investigated whether CarboTaxol promoted better activation of DC, professional antigen-presenting cells playing a major role in T-cell priming. CarboTaxol did not promote the expansion of spleen DC subsets such as conventional (c) DC1, DC2 and monocyte-derived DC (mDC) in tumor-free mice (Supplemental Figure 5 A-B). A trend towards an increased proportion of CD80+ cells in the cDC1 and cDC2 population was observed in CarboTaxol treated groups although expansion of absolute cell counts of CD80+ cDC1 cells or CD80+ cDC2 cells was not observed (Supplemental Figure 5 C-F). Moreover, the serum level of inflammatory cytokines IFNα and IL-1β were not impacted by CarboTaxol 4 days after treatment (Supplemental Figure 5G). Altogether, these results do not support an effect of CarboTaxol on DC activation.

### CarboTaxol induces TCF1+CD8+ T cells in a tumor-independent manner and promotes stem-like TCF1+PD-1+CD8+ T-cell infiltration in the TME

We then evaluated the phenotype of CD8+ T cells during priming in tumor-free mice according to the scheme shown in Figure 5A. We observed an increased TCF1 expression in CD8+ T cells in the spleen (Figure 5B) and in the blood (Figure 5C) of tumor free mice treated with CarboTaxol as compared to untreated mice. Notably, besides an increased mean fluorescence intensity (MFI), we observed a significant expansion of TCF1+CD8+ T cells in the spleen and blood after CarboTaxol treatment, independent of ChAdOx1 vaccination (Figure 5 D-E). Antigen-experienced CD44+TCF1+ CD8+ T cells were also expanded after CarboTaxol treatment compared to the control or vaccines only groups (Figure 5 D-E). Of note, we did not observe depletion of effector or central memory CD8+ T-cell subsets in the spleen and blood after CarboTaxol (Supplemental Figure 6 A-C), confirming that the expansion of CD8+ TCF1+ T cells was not conjugated with the depletion of effector or central memory cells, but rather seemed to correlate with an expansion of total CD8+ T cells in the spleen (supplemental Figure 6B). Because TCF1 expression is known to promote better CD8+ T cell proliferation and activation ^18,19^, these data suggested that the adjuvant effect of CarboTaxol might result from the increase in TCF1 expression.

**Figure 5:**
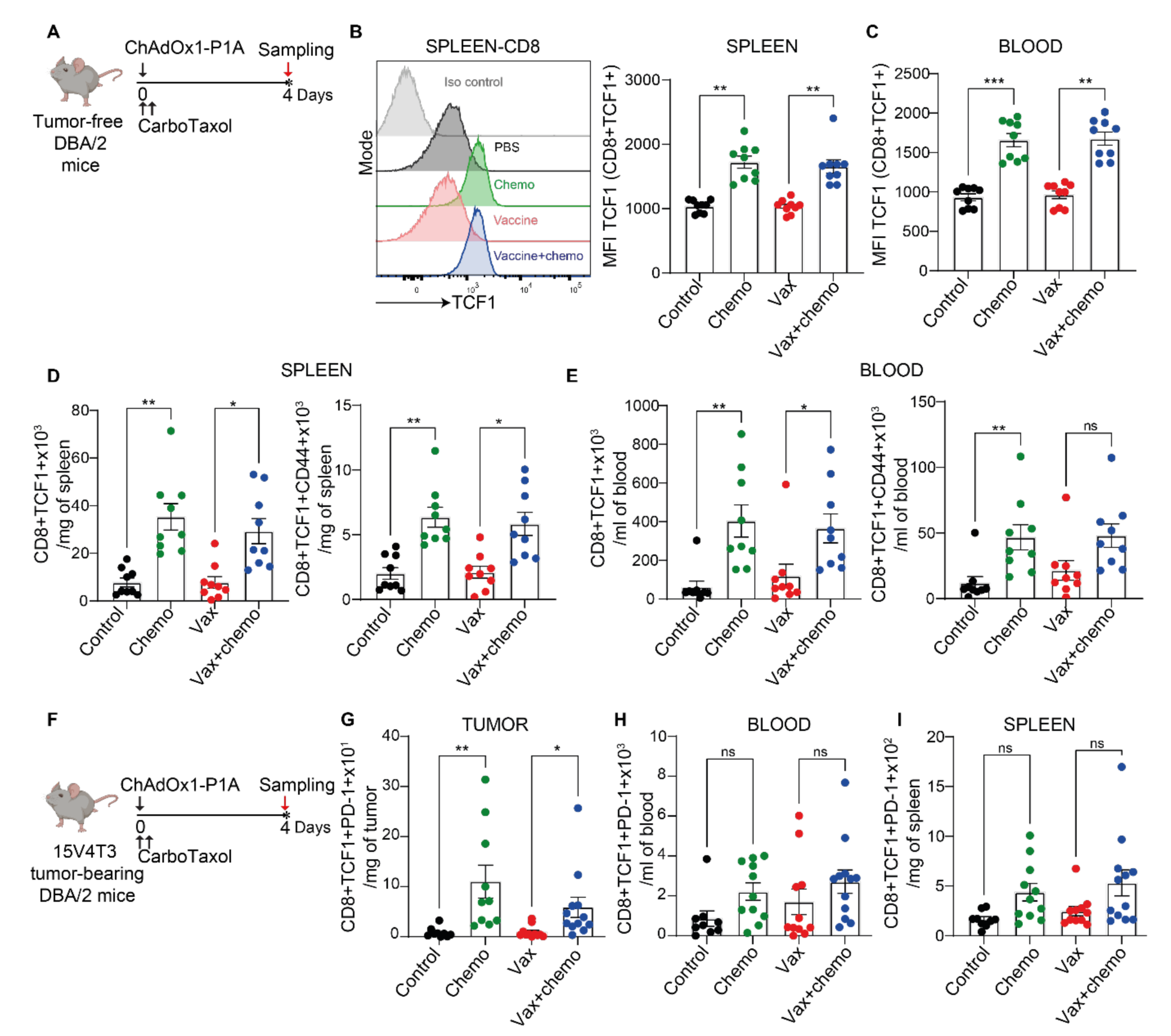
CarboTaxol induces TCF1+CD8+ T cells in a tumor-independent manner and promotes stem-like TCF1+PD-1+CD8+ T-cell infiltration in the TME. **(A)** Schematic representing timeline of experiment. Tumor free DBA/2 mice were vaccinated with ChAdOx1- P1A or PBS sham and treated with CarboTaxol on the same day. On day 4 post treatment, spleen and blood were harvested and CD8+ T cell phenotype was investigated by flow cytometry. **(B)** The MFI level of TCF1 in CD8+ T cells is shown in the spleen of the different groups. **(C)** The MFI of TCF1 in CD8+ T cells in the blood is shown. (**D-E**) The numbers of CD8+ TCF1+ T cells and CD44+TCF1+CD8+ T cells are shown in the spleen (**D**) and blood (**E**). The data are pooled from 2 independent experiments and mean± SEM for each group is shown. Each symbol represents an individual mouse (n=9 mice per group). (**F**) DBA/2 mice were implanted with 15V4T3 cells and 14 days after tumor implantation, mice were vaccinated with ChAdOx1-P1A or PBS sham and treated with CarboTaxol. (**G-I**) The TCF1+ PD-1+ CD8+ T cell numbers are shown in the TME (**G**), the spleen (**H**), and blood (**I**). The data are representative of 3 independent experiments and mean ± SEM for each group is shown. Each symbol represents an individual mouse (n=9 to 12 mice per group). Statistically significant differences between groups were determined by a Kruskal-Wallis test with Dunn’s multiple comparisons test. ns non-significant, * p ≤ 0.05, ** p ≤ 0.01, *** p ≤ 0.001.

We further examined whether the effect of CarboTaxol on TCF1 extended to tumor-bearing mice. Fourteen days after tumor implantation, we vaccinated DBA/2 mice with or without CarboTaxol, and analyzed spleen, blood and tumor samples 4 days later (Figure 5F). TCF1 expression as measured by MFI was increased in spleen and blood CD8+ T cells (Supplemental Figure 6 D-F). Moreover, the TCF1+CD8+ and TCF1+CD44+CD8+ T cell populations were expanded in the spleen and tumor, (Supplemental Figure 6 G-H). Importantly, a significant increase in TCF1+PD-1+ CD8+ T cells in the tumor was also observed (Figure 5G), with a similar trend detected in blood and spleen (Figure 5 H-I). Furthermore, tumor samples collected after triple combination of CarboTaxol, ChAdOx1/MVA and anti-PD-1 exhibited the highest proportion and number of stem-like TCF1+PD-1+CD8+ T cells (Supplemental Figure 6 I-J). These cells are known as stem-like progenitor exhausted T cells and are often enriched in the TME and tumor-draining lymph nodes ^10^. They display enhanced functionality and proliferation capacity in the TME compared to terminally differentiated TCF1-PD-1+CD8+ T cells ^20^. Moreover, stem-like progenitor exhausted CD8+ T cells respond better to anti-PD-1 and appear to be the main population driving tumor rejection upon PD-1 blockade. Accordingly, they have been correlated with improved anti-PD-1 treatment response ^10^. Therefore, this early expansion of TCF1+PD-1+CD8+ T cells among tumor-infiltrating lymphocytes after CarboTaxol could explain the superior tumor control we observed in the triple combination setting with CarboTaxol, vaccines and anti-PD-1.

### The immune adjuvant effect of CarboTaxol depends on the TCF1/β-catenin pathway

TCF1 is a transcription factor that includes several isoforms, with the long isoform able to bind to β-catenin, which is necessary for its transcriptional activity ^11^. TCF1 was shown to improve newly activated CD8+T cell proliferation *in vitro* in a β-catenin dependent manner ^19^. Hence, we asked whether TCF1 and its β-catenin-dependent-transcriptional activity were responsible for the adjuvant effect of CarboTaxol. To test this hypothesis, we employed a small molecule inhibitor, iCRT3, which can block TCF1 binding to β-catenin ^11^. We administered iCRT3 to tumor-free DBA/2 mice daily for 6 days starting 1 day before vaccination and CarboTaxol (Figure 6A). Four days after vaccination, we investigated the transcriptional activity of TCF1 by evaluating the expression of Ly108 (SLAMF6 in human), which is a direct target of the β- catenin/TCF1 pathway and has previously been used to assess TCF1 transcriptional activity ^21^. Using flow cytometry, we observed no change in TCF1 MFI between iCRT3 and untreated groups (Figure 6B). However, we detected a reduced expression of Ly108 in TCF1+ CD8+T cells from iCRT3-treated group, indicating the inhibitor was functional (Figure 6B). The effect of iCRT3 was further confirmed by a decreased expression of BCL6, another TCF1 transcriptional target, which is linked to the maintenance of progenitor exhausted stem-like cells ^22^ (Supplemental Figure 7 A-B). Remarkably, the addition of iCRT3 to CarboTaxol and the vaccines abrogated the adjuvant effect of CarboTaxol post-prime, as the proportion of CD8+IFNγ+ T cells and CD8+TNFα+ T cells 15 days after the prime was decreased as compared to the vaccines + CarboTaxol group (Figure 6C). A similar trend was detected 14 days after the boost (Figure 6D).

**Figure 6:**
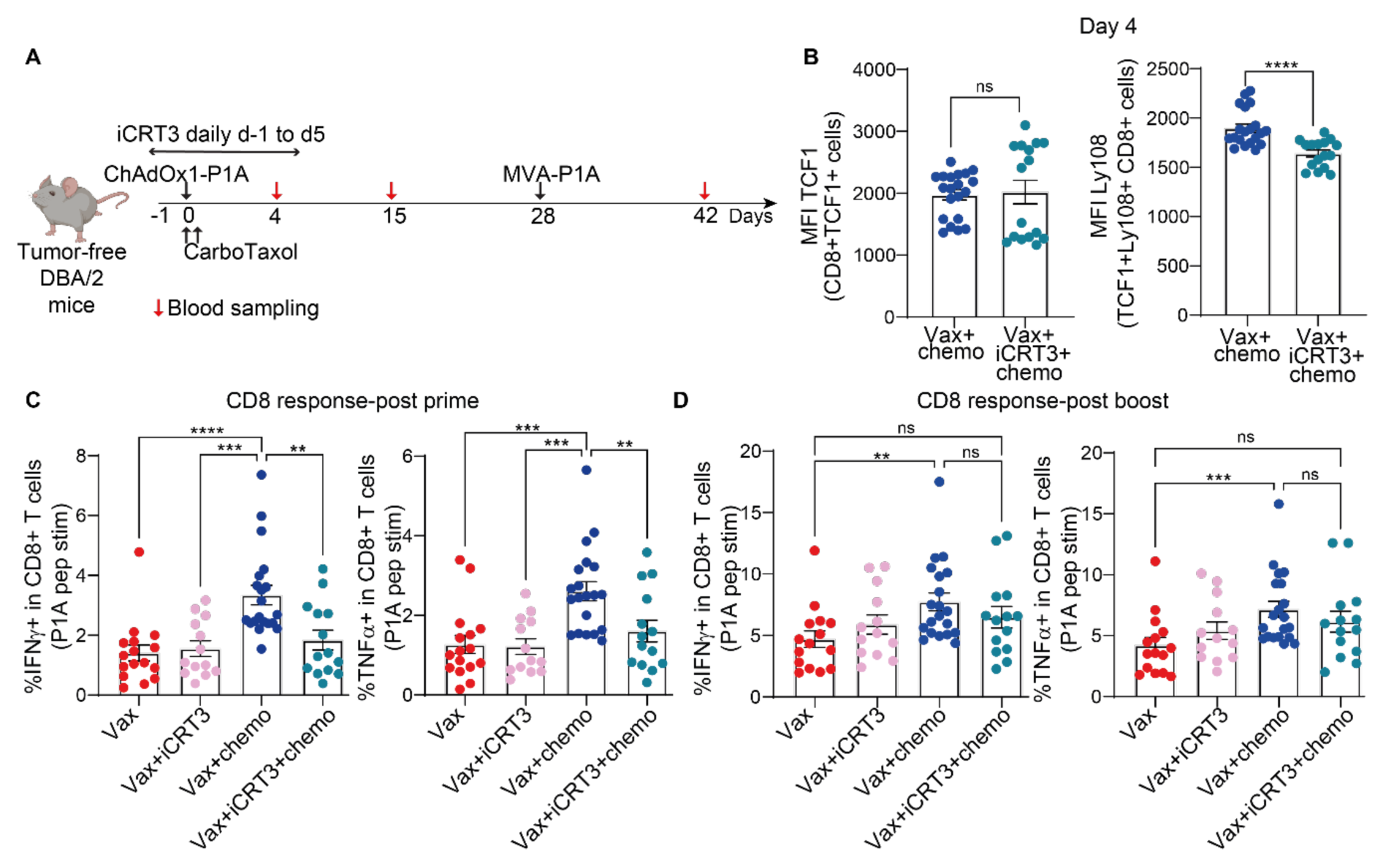
The adjuvant effect of CarboTaxol depends on the TCF1/β-catenin pathway. **(A)** Schematic representing timeline of experiment. Tumor free DBA/2 mice were vaccinated with ChAdOx1- P1A or PBS sham and treated with CarboTaxol on the same day. Additionally, some mice were treated with iCRT3 daily for 6 days starting one day prior to the prime vaccination. **(B)** At day 4 post-prime, blood was harvested and TCF1 expression as well as its transcriptional target Ly108 (SLAMF6 in human) were investigated by flow cytometry. The MFI level of TCF1 in TCF1+CD8+ T cells and the MFI level of Ly108 in TCF1+Ly108+CD8+ T cells is shown as mean ± SEM. (**C-D**) The proportion of IFNγ-producing and TNFα-producing P1A-specific CD8+ T cells are shown as mean ± SEM post-prime (day 15) (**C**) and post-boost (day 42) (**D**). Data are pooled from at least 2 independent experiments (n= 13 to 20 mice). Statistically significant differences between groups were determined by a Mann-Whitney comparisons test. ns non-significant, * p ≤ 0.05, ** p ≤ 0.01, *** p ≤ 0.001.

To further confirm the role of TCF1 expression in CD8+ T cells as a mechanism for the vaccine adjuvant effect of chemotherapy, we also evaluated the impact of other chemotherapy drugs on the ChAdOx1-P1A-induced CD8+ T cell-response and TCF1+CD8+ T- cell expansion in tumor-free mice (Supplemental Figure 7C). On day 4 post treatment, cyclophosphamide promoted TCF1+CD8+ T-cell expansion (supplemental Figure 7D), and the proportion of TCF1+ cells in CD8+ T cells positively correlated with the P1A-specific CD8+ T- cell response induced by ChAdOx1-P1A as measured on day 15 (Supplemental Figure 7E). In contrast, gemcitabine, which did not promote TCF1 expression to the same extent as cyclophosphamide or CarboTaxol, did not show an immune adjuvant effect (Supplemental Figure 7D-E). These results support the role of TCF1 expression in the adjuvant effect of chemotherapy.

### CarboTaxol expands *TCF7+CD8+* T cells in ovarian cancer patients

To better understand the human translational relevance of our findings, we utilized RNA sequencing data from a longitudinal study investigating immune functions in patients with advanced epithelial ovarian cancer to explore TCF7 (the gene encoding TCF1) expression in CD8+ T cells ^23^. These patients underwent SoC neoadjuvant carboplatin and paclitaxel chemotherapy, with the inclusion of dexamethasone for prophylaxis of paclitaxel-associated hypersensitivity reactions. Single-cell RNA sequencing of the patients’ PBMCs was performed at baseline and after the third cycle (C3) of carboplatin and paclitaxel treatment (Figure 7A). We compared the proportion of *CD8*+ T cells expressing *TCF7* as well as *PDCD1* (PD-1 gene). Though the proportion of *CD8*+ T cells remained similar (Figure 7B), the proportion of *TCF7*+*CD8*+ T cells was increased after the third cycle of carboplatin and paclitaxel treatment compared to the baseline (Figure 7C). A trend of a slight increase in *TCF7*+*PDCD1*+*CD8*+ T cells was also detected (Figure 7D). Furthermore, the average expression level of *TCF7* in *CD8*+ T cells was increased at C3 compared to the baseline (Figure 7E), indicating a positive impact of carboplatin and paclitaxel on *TCF7* expression in *CD8*+ T cells from patients with epithelial ovarian cancer. These data confirm in the human setting our observation of an expansion of TCF1+CD8+ T cells after CarboTaxol treatment.

**Figure 7:**
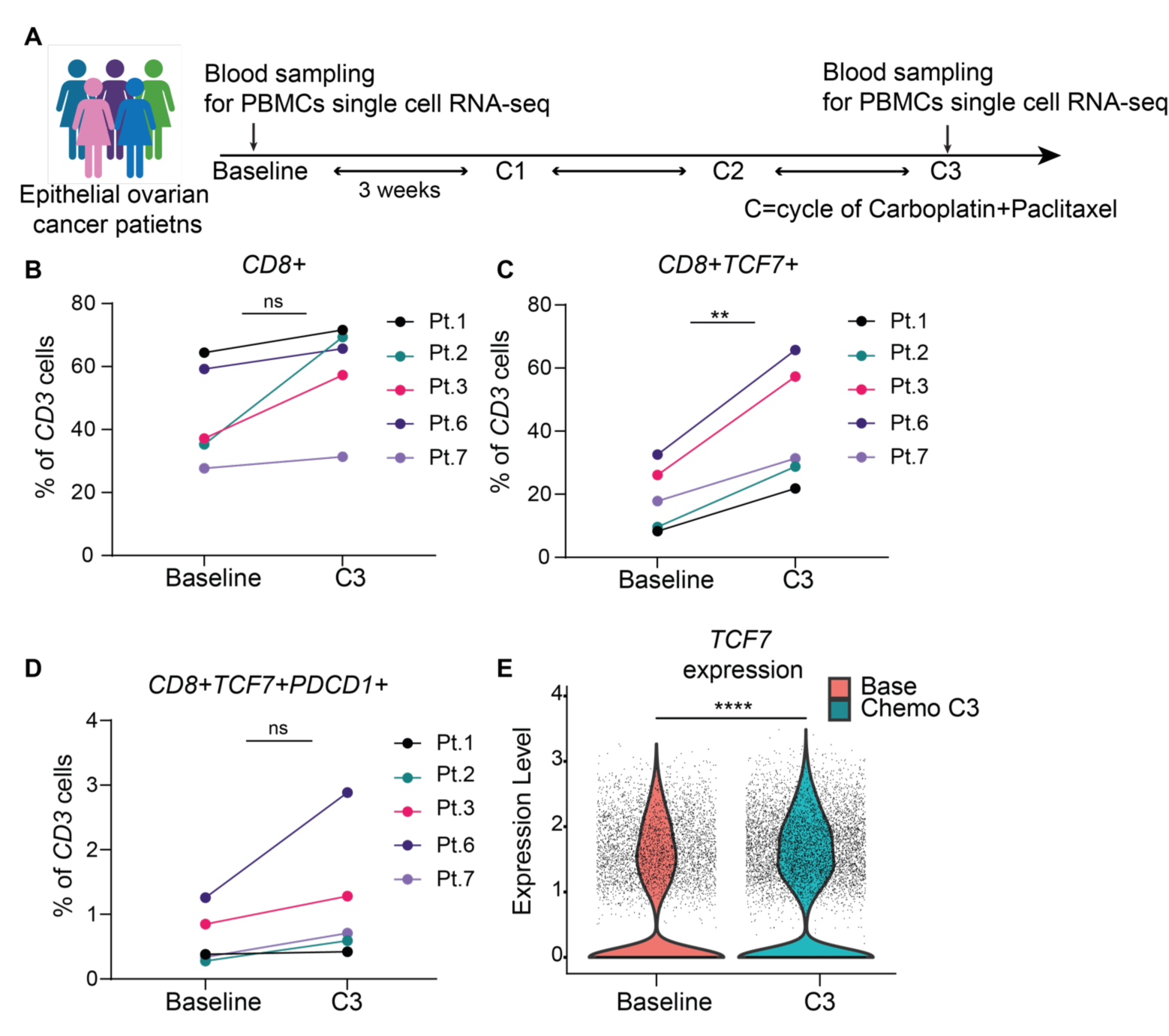
CarboTaxol expands *TCF7+CD8+* T cells in ovarian cancer patients. (**A**) The scheme was adapted from *Liu et al.,2022.* This longitudinal study performed single cell RNA-sequencing using patients’ blood, collected at baseline and after 3 cycles of chemotherapy (C3). (**B-D**) The proportion of *CD8*+ T cells (**B**), of *TCF7*+ *CD8+* T cells (**C**) and of *TCF7+ PDCD1+ CD8+* T cells (**D**) among *CD3*+ cells between at baseline and C3 are shown as mean± SEM. After testing for normality, statistically significant differences between groups were determined by a Paired t test. (**E**) Expression level of *TCF7* in *CD8*+ cells is shown at baseline and at C3. Statistically significant differences between groups were determined by one-sided Wilcoxon rank sum test. ns non-significant, ** p ≤ 0.01, **** p ≤ 0.0001.

## Discussion

In the clinic, therapeutic cancer vaccines will likely be given in combination with SoC, which often comprises chemotherapy and checkpoint inhibitors. Yet, given their different mechanisms of action, it is unclear to what extent these modalities can be synergistic, and how best to combine them, particularly with regard to chemotherapy and its cytotoxic effects. While some chemotherapies have immunomodulatory properties, how to harness these properties for the benefit of vaccines or immunotherapies remains unclear and may depend on a precise schedule of administration ^5^. Here, we explored the combination of CarboTaxol with viral vector vaccines ChAdOx1/MVA expressing the murine MAGE-type antigen P1A, given with or without anti-PD-1. We observed that CarboTaxol increased the magnitude of the vaccine-induced P1A-specific CD8+ T cells, highlighting an immunomodulatory effect of CarboTaxol. This adjuvant effect was tumor-independent and observed with different vaccine platforms. The immunomodulatory role of CarboTaxol depended on its administration timing relative to the vaccines, similar to other chemotherapies tested in combination with immunotherapies, such as doxorubicin and cyclophosphamide ^24–26^. Here, CarboTaxol needed to be given on the same day as the prime or boost vaccination to have an adjuvant effect, implying a role of CarboTaxol on the priming environment during vaccination. Notably, cyclophosphamide, but not gemcitabine, displayed a similar immune adjuvant effect, indicating that this effect might also depend on the class of chemotherapy.

For some cancer types, the SoC also includes immune checkpoint blockade, such as anti-PD-1, which aims to prevent T cell exhaustion of tumor-infiltrating lymphocytes. In the vaccination setting, it is therefore particularly relevant to include checkpoint blockade in the therapeutic strategy. Our previous publication showed that ChAdOx1/MVA-P1A synergized with anti-PD-1, promoting superior tumor control and survival as compared to vaccines alone^3^. The triple combination of ChAdOx1/MVA, anti-PD-1 and CarboTaxol outperformed the relevant double therapy combination groups, displaying the best tumor growth control, tumor clearance and survival in 15V4T3 and TC-1 tumor models. Furthermore, CarboTaxol extended the length of the possible administration window of anti-PD-1 treatment relative to cancer vaccines. Indeed, the timing of administration of anti-PD-1 was previously shown to be crucial for the synergistic effect with vaccination in the TC-1 tumor model ^15^. Anti-PD-1 given prior to vaccination resulted in the loss of treatment synergy compared to anti-PD-1 given after vaccination, which was also confirmed in the 15V4T3 tumor model in this study. Strikingly, CarboTaxol combined with the vaccines could rescue the negative effect of anti-PD-1 given prior to vaccination, indicating the importance of CarboTaxol in driving the synergy behind this triple combination therapy, and supporting a schedule in which the SoC is started without delay while the vaccine is being prepared or the patient is checked for vaccine eligibility. Corroborating this finding, NSCLC patients resistant to ICB and platinum-based chemotherapy, and subsequently treated with a peptide vaccine displayed significantly improved overall median survival compared to chemotherapy alone ^27^.

We explored the mechanism of action of this immune adjuvant effect of CarboTaxol. Previous studies reported that CarboTaxol depletes CD11b+Gr1+ cells, and that vaccination during myeloid cell depletion promotes better tumor growth control ^6,7^. On day 4 after CarboTaxol, we also observed a depletion of CD11b+Gr1^hi^Ly6G+ cells. However, we could not link this depletion to the synergistic effects of CarboTaxol with vaccines in the 15V4T3 tumor model. Depletion of CD11b+Ly6G+ cells with anti-Gr1 or anti-Ly6G did not reproduce the impact of CarboTaxol on tumor growth control, survival, and immunogenicity when combined with ChAdOx1/MVA-P1A vaccination, suggesting there is another underlying mechanism behind the synergy of CarboTaxol with ChAdOx1/MVA vaccines. The fact that the immune adjuvant effect of CarboTaxol was also observed in tumor-free mice also excluded ICD as the mechanism involved. We did not observe clear evidence for an effect on DC activation or maturation, and we therefore investigated the phenotype of CD8+ T cells. Remarkably, CarboTaxol increased TCF1 expression in CD8+ T cells 4 days after treatment and expanded stem-like TCF1+CD8+ T cells. Moreover, CarboTaxol promoted the recruitment of progenitor exhausted stem-like TCF1+PD-1+CD8+ T cells in the TME. The expression of TCF1 in T cells confers a better activation, proliferation, and differentiation potential as compared to TCF1 negative cells ^18,19,28^. These populations were correlated with stronger anti-tumor responses as well as improved responsiveness to anti-PD-1 treatment ^10,29,30^. Therefore, TCF1 plays a pivotal role in promoting CD8+ T cell fitness when confronted to the immunosuppressive TME, providing a potential explanation for the enhanced tumor control observed in mice treated with vaccines and CarboTaxol or with the triple combination strategy. Moreover, we showed for the first time that TCF1 transcriptional activity during priming promotes an immune adjuvant effect for cancer vaccines. Indeed, the adjuvant effect of CarboTaxol was abrogated by iCRT3, an inhibitor of TCF1/β-catenin interaction. These data are in line with a recent publication showing that TCF1 expression is necessary for the establishment of an immune response against poorly immunogenic tumors ^20^. Altogether, TCF1 expression in T cells following CarboTaxol treatment is beneficial to raise an efficient anti-tumor response and can be used to potentiate cancer vaccine efficacy.

The mechanism by which CarboTaxol promotes TCF1 expression in T cells has yet to be understood. Paclitaxel was shown to activate the WNT/β-catenin pathway ^31^, which is known to regulate TCF1 expression ^32,33^. Given our results showing the importance of the TCF1 and β-catenin interaction, it is possible that CarboTaxol impacts the WNT/β-catenin upstream of TCF1, promoting TCF1 expression. However, TCF1 regulation is complex and can be modulated by different pathways ^34–36^. Notably, both carboplatin and paclitaxel were shown to induce the cGas/STING pathway ^37,38^, which acts upstream of TCF1 and promotes its expression and maintenance in CD8+ T cells in a cell-autonomous way ^36^. Although we did not observe a change in serum level of IFNα after CarboTaxol treatment, the impact of CarboTaxol on cGAS/STING cannot be excluded based on the data presented here.

Importantly, we observed an increase in the proportion of *CD8*+*TCF7*+ cells among *CD3*+ T cells in the blood of ovarian cancer patients treated with three cycles of CarboTaxol. This corroborates our findings from the mouse studies that CarboTaxol can indeed expand stem-like TCF1+CD8+ T cells. Despite the small cohort size and the fact that patients were additionally treated with the immunosuppressive drug dexamethasone ^23^, the effect of CarboTaxol on *TCF7* expression in *CD8*+ T cells could be detected, suggesting a consistent effect of CarboTaxol. Collectively, these data hold substantial promise for the clinical testing of the combination of cancer vaccines with SoC that comprises chemotherapy and anti-PD-1.

### Limitations of the study

There are some limitations to the studies presented herein. The studies were performed primarily with one tumor model, the 15V4T3 mastocytoma, with P1A as the tumor antigen. However, the improvement of tumor control and survival by the triple combination therapy was confirmed in a second tumor model, the lung carcinoma TC1, using the HPV antigens E6 and E7 as tumor antigens. One outstanding question in this study is how to translate the chemotherapy dose and schedule from a mouse model to cancer patients. Different chemotherapies can have a range of immunomodulatory effects, but these effects are dependent on the dose and timing of administration. Therefore, it remains challenging to predict the immunomodulatory effect of a given chemotherapy in patients, often treated at the maximum tolerated dose (MTD) for several cycles. However, it is worth noting that ovarian cancer patients treated at the MTD of CarboTaxol still revealed an increased *TCF7* expression at transcriptional level after 3 cycles of carboTaxol treatment. Further analysis of TCF1 expression at the protein level across various time points during chemotherapy, both with and without immunotherapy, will provide valuable data to confirm translatability to humans.

## Methods Lead contact

Further information and requests for resources and reagents should be directed to and will be fulfilled by lead author, Carol Leung.

## Materials availability

Requests for ChAdOx1 and MVA viral vector vaccines can be directed to Barinthus Biotherapeutics Ltd.

## Data and code availability

Any additional information required to reanalyze the data reported in this paper is available from the lead contact upon request.

## Mice

Eight- to eleven-week-old female C57BL/6 and DBA/2 mice used in this study were purchase from Envigo, UK. The studies performed here were carried out in accordance with the terms of the UK Animals Act Project License (PPL) P0D369534.

## Vaccines

ChAdOx1 and MVA viral vectors encoding P1A were cloned and produced as previously described ^3^. For the HPV E6/E7 vaccines, the coding sequence of the E6 and E7 CD8 epitope peptides (E6-CKQQLLRREVYDFAFRDLCIVYRDG; E7-GQAEPDRAHYNIVTFCCKCD) were purchased as strings DNA fragments from GeneArt (Thermo Fisher Scientific), then inserted in the ChAdOx1 and MVA vectors as previously described ^39^. A sequence coding for the 26 amino acid transmembrane domain of the MHC-II invariant chain (GALYTGVSVLVALLLAGQATTAYFLY) was linked to the N terminus of the transgene to improve immunogenicity in the ChAdOx1 vector. The purity and identity of the viral vectors were confirmed by PCR. The SNP P1A vaccine was produced in Dr. Robert Seder’s lab in NIH with an established protocol ^40^, in which the P1A peptides were conjugated with the hydrophobic block containing a TLR-7/8 agonist.

### Tumor cell cultures and tumor implantation

15V4T3 tumor cell line is a mastocytoma line of DBA/2 origin and expresses P1A ^41^. TC1 tumor cell line, which is of C57BL/6 origin, expresses the E6/E7 antigens and was a kind gift from Dr. Robert Seder’s lab in NIH.

To initiate tumor growth, 1x10^6^ 15V4T3 cells or 1x10^5^ TC1 cells were injected subcutaneously (s.c.) in the right flank of the mice. Upon the development of palpable tumors, measurements of tumor length and width were carried out 2-3 times per week using digital calipers. The volume of tumor masses (V) was calculated according to the formula: V = ((length (mm) x width^2^ (mm)) x 0.52). On average, tumor studies were started when tumor size reached around 50 mm^3^. Mice were sacrificed via cervical dislocation when tumor size reached 15mm in any direction.

## Vaccination

Vaccinations were performed under inhalational anesthesia, using isoflurane as the anesthetic agent. Vaccines were administrated via intramuscular injection (i.m.) into the right thigh at either a high or low dose regimen. The high dose regimen comprised of 10^8^ (high) infectious units (IU) of ChAdOx1 virus and 10^7^ (high) plaque forming units (PFU) of MVA virus. This dose was given as part of a 4-week apart regime and was used for immunogenicity study in tumor-free mice. Otherwise, the low dose comprised of 10^7^ IU of ChAdOx1 and 10^6^ PFU of MVA was used. Vaccines were diluted in sterile PBS for injections and administered in a total volume of 50µl per animal. To control for vaccination, ChAdOx1 and MVA expressing DPY, an irrelevant human protein, or PBS were used. The SNP P1A vaccine was prepared in sterile PBS and administered subcutaneously.

### Administration of monoclonal antibody, chemotherapy and iCRT3

Mice were treated with monoclonal antibody or the relevant isotype control via intraperitoneal (i.p) injections. 100 μg of anti-PD-1 (RMPI-14, BioXcell) antibody was administrated at a frequency of 1 injection every third day for a total of 3 to 4 injections. Anti- Gr1 or anti-Ly6G antibodies (RB6-8C5 and 1A8 respectively, BioXcell) were administered at a dose of 200 µg twice weekly for a total of 5 injections. Chemotherapy drugs were dissolved in 200 μl sodium chloride solution and administered via i.p injections. Carboplatin was given in 1 dose of 40 mg/kg and paclitaxel was given in 2 doses of 20 mg/kg in 24 hours (h) apart. Cyclophosphamide was given in 1 dose of 50 mg/kg and Gemcitabine was given at 1 dose of 60 mg/kg. For administering iCRT3, mice were injected with a dose of 20 mg/kg (and 40mg/kg on day 3) daily for 6 days via i.p. iCRT3 was diluted in a total of 150 μl of PEG300 (45%), tween 80 (5%) and sodium chloride (50 %).

### Stimulation assay for antigen-specific T-cell responses

PBMCs, tumor, spleen and bone marrow cell suspension were obtained as previously described ^42,43^, and stimulated with αCD28 (2 μg/ml, Tonbo Biosciences) and DNAse I (20 μg/ml, Roche) and P1A peptide pool or E6/E7 pool (4 μg/ml, Peptides and Elephant) at 37°C 5% CO2 for 5 hours, with the addition of Brefeldin A (10 μg/ml, BioLegend) after 1 hour to promote intracellular cytokine accumulation.

### Flow cytometry staining

Cell suspension from tumor, spleen and blood were kept on ice during processing and all incubations and centrifugations were performed at 4°C. Blocking step using 50 µl Fc block, anti-mouse CD16/CD32 antibody (BD pharmingen) was performed prior to the staining. Then, cells were stained with viability dye and specific fluorophore-conjugated antibodies diluted in PBS for 25 to 30 minutes at 4°C. To assess the presence of cytokine production or cytoplasmic markers in T cells, the cells were fixed and permeabilized with the BD Cytofix/Cytoperm™ buffers (BD pharmingen) according to manufacturer’s instruction for 20min at 4°C and then stained for further intracellular proteins for 25 to 30 minutes at 4°C. Alternatively, to assess the presence of nuclear markers, such as TCF1, cells were fixed with eBioscience™ Foxp3 / Transcription Factor Staining Buffer Set (ebioscience) according to manufacturer’s instruction, for 35min at 4°C. For staining of nuclear proteins, and if included, cytoplasmic markers, antibodies were incubated for 12 to 16 h at 4°C. Details on the antibodies used can be found in Table 1. The next day, samples were acquired on a BD Fortessa cytometer and data were analyzed using FlowJo software. Of note, for experiments using anti-Gr1 antibody, cells were stained intracellularly for the Gr1, Ly6C and Ly6G markers as the epitope for these markers were masked by anti-Gr1 antibody, as shown previously ^44^.

**Table 1.**

| Marker | Clone | Company |
| --- | --- | --- |
| CD4 | GK1.5 | BioLegend |
| CD8a | 53-6.7 | BioLegend or BD Horizon |
| IFNγ | XMG1.2 | BioLegend |
| TNFα | MP6-XT22 | BioLegend |
| PD-1 | 29F.1A12 | BioLegend |
| Gr1 | RB6-8C5 | BioLegend |
| CD11b | M1/70 | BioLegend |
| CD11c | N418 | BioLegend |
| Ly6G | 1A8 | BioLegend |
| Ly6C | HK1.4 | BioLegend |
| CD38 | 90 | BioLegend |
| TCF1 | C63D9 | Cell Signaling technology |
| CD3 | 17A2 | BioLegend or BD Horizon |
| CD44 | IM7 | BioLegend |
| CD62L | MEL-14 | BioLegend |
| Ly108 | 13G3-19D | eBioscience |
| BCL6 | 7D1 | BioLegend |
| CD45 | 30-F11 | BioLegend |
| MHCII | 2G9 | BD Horizon |
| CD172a | P84 | BioLegend |
| XCR1 | ZET | eBioscience |
| CD64 | x54-5/7.1 | BioLegend |
| F4/80 | BM8 | BioLegend |
| CD80 | 16-10A1 | BioLegend |
| Live and Dead | (Fixable Viability Dye eFluor 506) | Ebioscience |

### Isolation of CD8+ T cells and Ly6G^+^ cells

Spleens from tumor-bearing DBA/2 mice (size range between 300 to 500 mm^3^) were collected. CD11b+Ly6G+ cells were extracted by magnetic selection with the Myeloid-Derived Suppressor Cell Isolation Kit, from Miltenyi Biotec following manufacturer’s instructions. Ly6G+ cells were then selected through LS column. The fraction of cells collected were the CD11b+GR1^hi^LY6G+ cells. However, for DBA/2 mice the purity was usually below 70% and this fraction contained many dead cells and debris. Therefore, the fraction of LY6G+ cells were then sorted to achieve a higher purity of 98 % or above. The cells were stained with Live/Dead, CD11b and Ly6G and then resuspended in 2 ml of FACS buffer for sorting. Ly6G+ cells were sorted on the Sony SH800z FACS sorter. For CD8+ T cells, spleens from naïve mice were collected and CD8+ T cells were isolated using negative selection with the CD8a+ T Cell Isolation Kit from Miltenyi Biotec following the manufacturer’s instructions.

### Proliferation assay

Following CD8+ T cell isolation, cells were stained with CellTrace Violet (5 μM). 1x10^5^ CD8+ T cells were plated in the 96-well round-bottom coated with 1 μg/ml of anti-CD3. CD11b+ Ly6G+ were added to the CD8+ T cell at different ratios. Anti-CD28 was added at a final concentration of 1 μg/ml into the plate. CD8+ T cell proliferation was assessed 72h later.

### ELISAs

IFNα and IL1β ELISAs were performed following manufacturer’s protocol (BioLegend). Briefly, one day prior to running the ELISA, 96-wells Nunc™ MaxiSorp™ ELISA plates were coated with 100μl of the relevant capture antibody. The next day, plates were blocked for 2 h before the addition of standard dilutions and samples to the appropriate wells. Samples were incubated at room temperature for 2 h with shaking. Then, samples were incubated with detection antibody for 1 h. Subsequently, avidin-HRP solution was added to each well for 20-30 minutes. Finally, 100 μL of TMB substrate solution were added and incubated in the dark for around 30 minutes before stopping the reaction with 100 μl of HCl. Absorbance was read with the CLARIOstar at 450 nm within 15 minutes.

### Single cell RNA-sequencing analysis

Single-cell RNA-seq data of PBMCs from ovarian cancer were downloaded from GEO (GSE164378- phs002862.v1.p1) based on a prior published study from Liu *et al* ^23^. The raw counts matrix was loaded into the Seurat package and combined with the metadata available.

The Seurat function PercentageFeatureSet was used to calculate the mitochondrial QC metrics. Cells with mitochondrial counts below 5% and unique feature counts between a user set range of 200 and 2500 were filtered for downstream analysis. The data was then log normalized with a scale factor of 10000 and the features that exhibit high cell-to-cell variation were identified. PCA was performed on the scaled data and cells were clustered using the Louvain algorithm. A UMAP embedding was used to visualize the cell clusters and broad cell types were annotated. The *CD3E* positive T cells from Seurat object was further subset into *CD8* and *CD4* for downstream analysis. The expression of *TCF7* and *PDCD1* were investigated in these broad cell types across baseline and chemo treated patients. The *CD8* object were re-clustered and visualized with a UMAP embedding. Differential expression of *TCF7* and *PDCD1* across baseline and chemo treated patients was investigated amongst the resulting cell types and visualized with a violin plot. Expression of genes were computed using one-sided Wilcoxon rank-sum test. Graphical representation of the proportion of the population of interest were made using Prism 8.0 software ((GraphPad software, CA, USA) and, after confirming the normality with a Shapiro-Wilk test, paired t test was performed to compare baseline and post chemotherapy proportions.

## Statistical analyses

Statistical analyses were performed using GraphPad Prism software (GraphPad software, CA, USA). Mann-Whitney test was performed to determine significant differences between the relevant groups regarding the P1A- specific immune responses induced by the vaccines, and the impact of each treatment addition on this response. To determine significant variation in cell populations and phenotypes across all groups, multiple comparisons were performed using Kruskal-Wallis test with Dunn’s multiple comparisons test. In murine tumor studies, a two-way ANOVA followed by Tukey’s post hoc test was used to determine statistically

significant differences in term of tumor growth between different groups. Survival curves for tumor studies were created using the Kaplan-Meier method and statistical significance in survival between different groups of mice was determined using the log-rank (Mantel-Cox) test. All P values < 0.05 were considered statistically significant.

## Study approval

All animal work was approved by the University of Oxford Animal Care and Ethical Review Committee and experimental procedures were carried out in accordance with the terms of the UK Animals (Scientific Procedures) Act Project License PB050649E.

## Acknowledgements

We thank V. Clark and H. Gray for animal husbandry, C. Farrow and K Dunning for critical manuscript reading. We thank Derin B Keskin and Panagiotis A Konstantinopoulos for sharing their single cell RNA-sequencing dataset published in “Improved T-cell Immunity Following Neoadjuvant Chemotherapy in Ovarian Cancer”. We thank Julia McCarthy and Svenja-Maria Pahlke for providing technical help in some harvest experiments. This work was supported by Ludwig Cancer Research. C.S.L. was supported by a fellowship from Swiss National Science Foundation (P300P3_155374).

## Author contributions

A.V.S.H., B.J.V.D.E., and C.S.L. conceived the experiments. L.N. and C.S.L. designed the experiments. L.N., A.W., V.P., J.M., E.S., S.P., and V.C. performed the experiments. R.R.V. designed and shared valuable resources. L.N., A.J., V.C., and C.S.L. analyzed the data. L.N. and C.S.L. wrote the manuscript, with all authors reviewed and edited the manuscript.

## Declaration of interests

A.V.S.H., B.J.V.D.E., and C.S.L. are inventors on a patent that covers viral vectors and methods for the prevention and treatment of cancer. A.V.S.H. is a co-founder of and shareholder in Barinthus Biotherapeutics Ltd which has supported the MAGE cancer vaccine program and has licensed rights to the ChAdOx1-MVA platform in cancer. B.J.V.D.E. is a scientific advisor in Barinthus Biotherapeutics Ltd. All other authors declare no conflict of interest.

