## Supplemental file for "Chemotherapy synergizes with cancer vaccines and expands stem-like TCF1+CD8+ T cells"

### Supplementary Materials

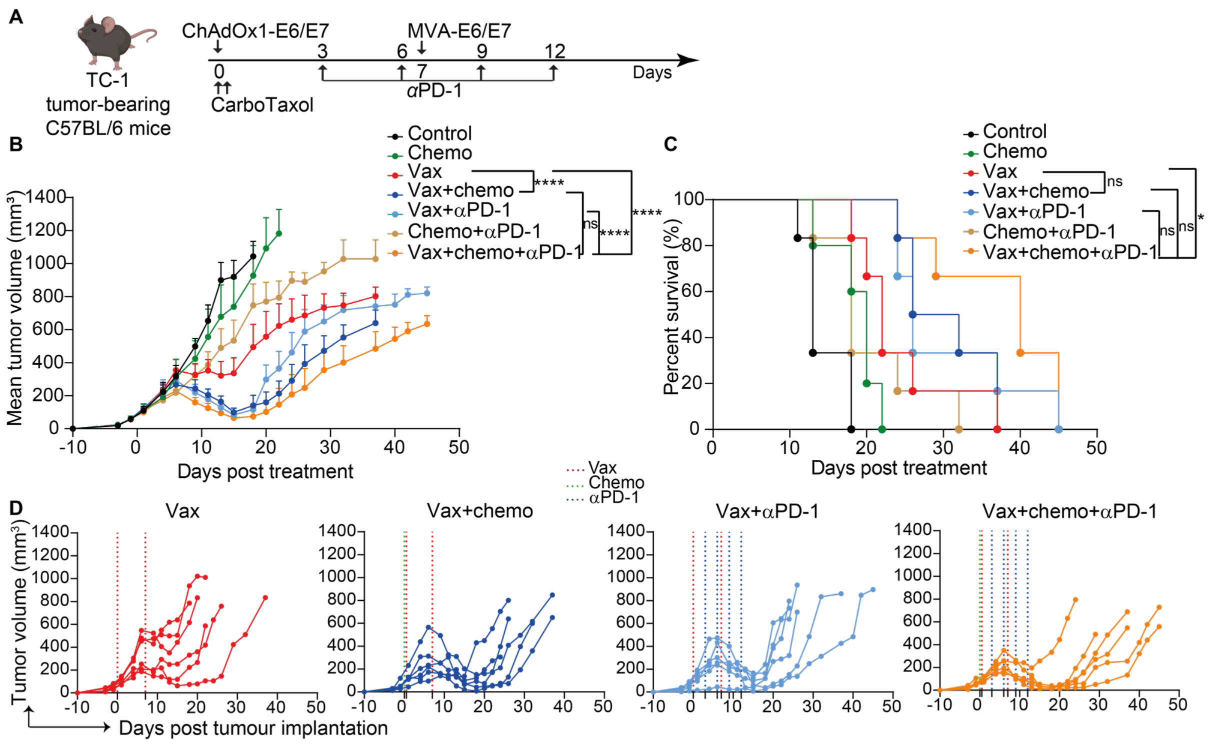

**Supplemental Figure 1: CarboTaxol, anti-PD1 and ChAdOx1/MVA expressing E6-E7 synergize for better tumor control and survival.**

(**A**) Schematic representing timeline of experiment. C57BL/6 mice were implanted with TC-1 cells to initiate tumor growth. 10 days after tumor implantation, mice were vaccinated with ChAdOx1-E6/E7 or ChAdOx1-DPY or PBS sham and treated with carboplatin and paclitaxel. 7 days later, the mice were vaccinated with MVA-E6/E7 or MVA-DPY and treated with anti-PD1 every third day for 4 injections.

(**B-D**) Mean tumor growth (**B**), mouse survival (**C**) and individual tumor growth curves (**D**) are shown in the indicated groups. Data are representative of 2 independent experiments (n=6 mice per groups). Tumor growth data in (B) are presented as mean tumor volume (mm$^3$) + SEM. Statistically significant differences in tumor volume between groups were determined by a two-way ANOVA followed by Tukey's post hoc test. Statistical differences in survival data (C) were determined by a Log Rank test. ns non-significant, * p ≤ 0.05 ****, p ≤ 0.0001.

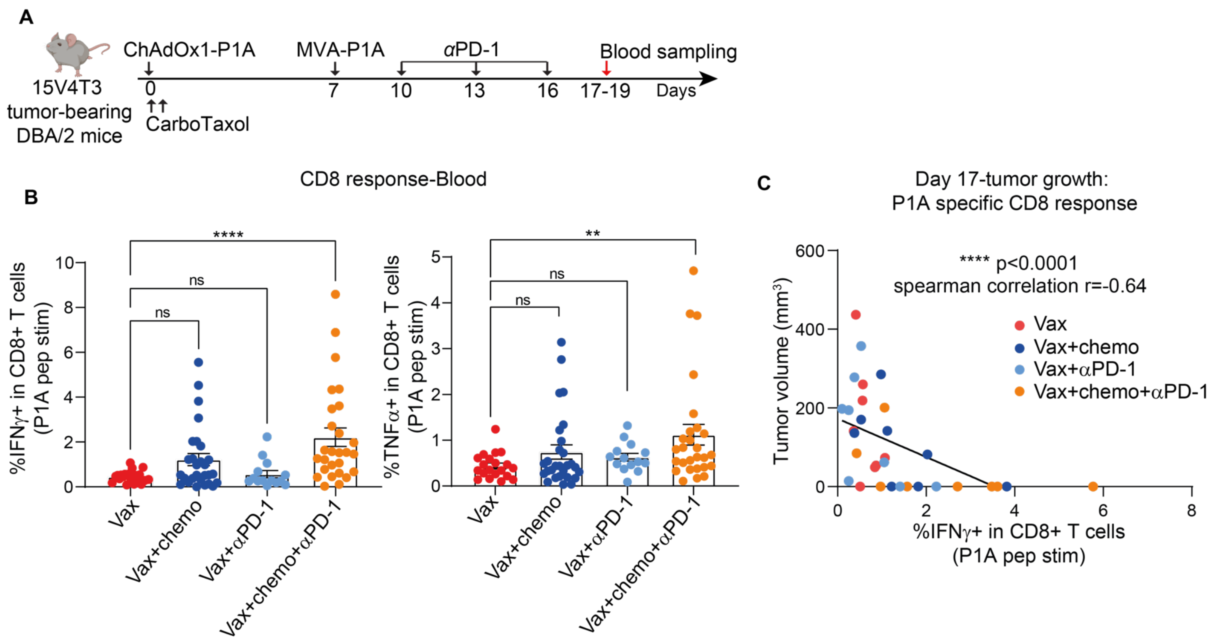


**Supplemental Figure 2: Anti-PD1 supports the P1A specific response long term.**
(**A**) Schematic representing timeline of experiment. DBA/2 mice were implanted with 15V4T3 cells and treated according to the scheme. Blood samples were collected 17 to 19 days after ChAdOx1-P1A and PBMCs were stimulated ex vivo with P1A peptides.
(**B**) Average of IFNγ+CD8+ response and TNFα+CD8+ response are shown. Data are pooled from at least 3 independent experiments and shown as the mean± SEM (n=14 to 28 mice per group). Each symbol represents an individual mouse. Statistically significant differences between groups were determined by Mann-Whitney test.
(**C**) Tumor size of day 17 post treatment was correlated with the magnitude of IFNγ producing P1A-specific CD8+ T-cells in the blood evaluated 17 days after prime ns for non-significant. Significance was determined by a Spearman Rank correlation test. ns non-significant, * $p \leq 0.05$, ** $p \leq 0.01$, *** $p \leq 0.001$, **** $p \leq 0.0001$.

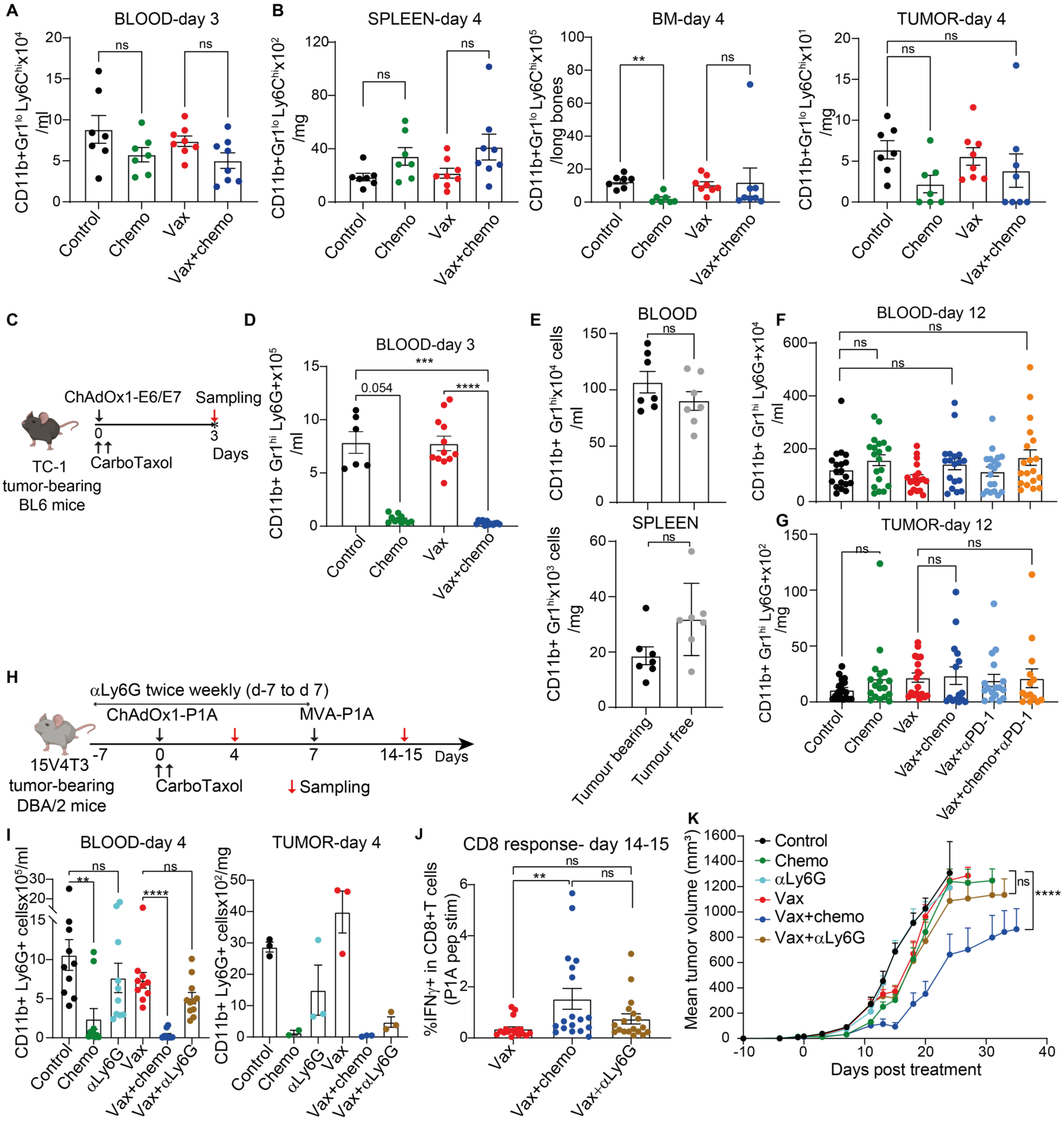
A
BLOOD-day 3
CD11b+Gr1lo Ly6Chi x10⁴ /ml
Control
Chemo
Vax
Vax+chemo
B
SPLEEN-day 4
CD11b+Gr1lo Ly6Chi x10² /mg
BM-day 4
CD11b+Gr1lo Ly6Chi x10⁵ /long bones
TUMOR-day 4
CD11b+Gr1lo Ly6Chi x10¹ /mg
C
ChAdOx1-E6/E7
Sampling
CarboTaxol
Days
TC-1 tumor-bearing BL6 mice
D
BLOOD-day 3
CD11b+ Gr1hi Ly6G+x10⁵ /ml
0.054
E
BLOOD
CD11b+ Gr1hi x10⁴ cells /ml
SPLEEN
CD11b+ Gr1hi x10³ cells /mg
Tumour bearing
Tumour free
F
BLOOD-day 12
CD11b+ Gr1hi Ly6G+x10⁴ /ml
G
TUMOR-day 12
CD11b+ Gr1hi Ly6G+x10² /mg
Vax+αPD-1
Vax+chemo+αPD-1
H
αLy6G twice weekly (d-7 to d 7)
ChAdOx1-P1A
MVA-P1A
14-15
Days
15V4T3 tumor-bearing DBA/2 mice
CarboTaxol
Sampling
I
BLOOD-day 4
CD11b+ Ly6G+ cellsx10⁵/ml
TUMOR-day 4
CD11b+ Ly6G+ cellsx10²/mg
αLy6G
Vax+αLy6G
J
CD8 response- day 14-15
%IFNγ+ in CD8+T cells (P1A pep stim)
K
Control
Chemo
αLy6G
Vax
Vax+chemo
Vax+αLy6G
Mean tumor volume (mm³)
Days post treatment

**Supplemental Figure 3: CarboTaxol does not impact CD11b+ Ly6C$^{hi}$ cells, potential M-MDSCs, and induces a transient CD11b+ Ly6G+ cell depletion.**

(**A-B**) DBA/2 mice implanted with 15V4T3 cells and vaccinated with ChAdOx1-P1A or PBS sham and treated with CarboTaxol. The numbers of CD11b+ GR1$^{lo}$ Ly6C$^{hi}$ are shown in the blood (**A**) spleen, BM, and tumor (**B**) at the indicated days.

(**C**) Schematic representing timeline of experiment. C57BL/6 mice were implanted with TC-1 cells to initiate tumor growth. 10 days after tumor implantation, mice were vaccinated with ChAdOx1-E6/E7 or ChAdOx1-DPY and treated with CarboTaxol.

(**D**) Numbers of CD11b+ GR1$^{hi}$ Ly6G+ are shown at day 4 post treatment. The data are representative of 2 independent experiments and mean ± SEM for each group are shown (n= 6 to 12 mice).

(**E**) The number of CD11b+GR1$^{hi}$ cells between non-treated tumor bearing at day 15 post tumour implantation with 15V4T3 and tumor free mice in the spleen and blood are shown as mean ± SEM. Each symbol represents an individual mouse (n=7 to 8 mice per group).

(**F-G**) CD11b+GR1$^{hi}$ cells at day 12 post treatment in the blood (**F**) and tumor (**G**) are shown. The data are pooled from 2 independent experiments and mean± SEM for each group is shown.

(**H**) Mice were treated with either CarboTaxol or with anti-Ly6G antibody twice a week starting 7 days prior to ChAdOx1-P1A vaccination. 7 days after the prime, mice were boosted with MVA-P1A.

(**I**) The numbers of CD11b+Ly6G+ cells in the tumor and blood are shown as mean ± SEM. Data are pooled from 2 independent experiments in the blood whilst tumor data show only one replicate.

Statistically significant differences between groups in (A-I) were determined by a Kruskal-Wallis test with Dunn's multiple comparisons test.

(**J**) The proportion of IFN$\gamma$+P1A specific CD8+ T cells are shown. Data were pooled from at least 2 independent experiments and mean ± SEM are shown (n= 9 to 17). Statistically significant differences were determined by Mann-Whitney test.

(**K**) Mean tumor growth (mm$^3$) + SEM is shown and is representative of 3 independent experiments. Statistically significant differences between groups were determined by two-way ANOVA followed by Tukey's post hoc test. ns non-significant, * $p \leq 0.05$, ** $p \leq 0.01$, *** $p \leq 0.001$, **** $p \leq 0.0001$.

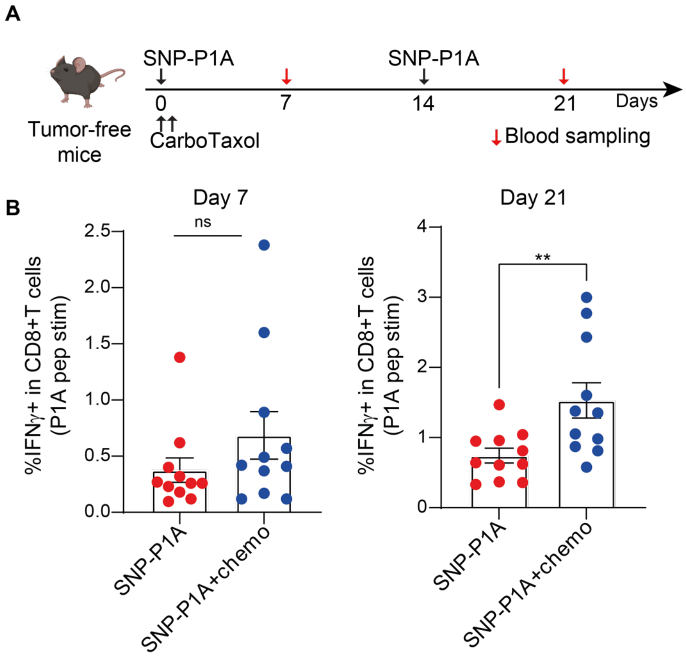


**Supplemental Figure 4: CarboTaxol acts as an adjuvant for SNP vaccine.**

(**A**) Schematic representing timeline of experiment. Mice were vaccinated with SNP-P1A either with or without CarboTaxol given with the prime. The response post prime and post boost were evaluated.

(**B**) Percentage of IFNγ-producing P1A specific CD8+ T cells is shown as mean± SEM. Each symbol represents an individual mouse and data are pooled from 2 independent experiments (n= 11 mice per groups). Statistically significant differences between groups were determined by a Mann-Whitney comparisons test. ns non-significant, ** $p \leq 0.01$.

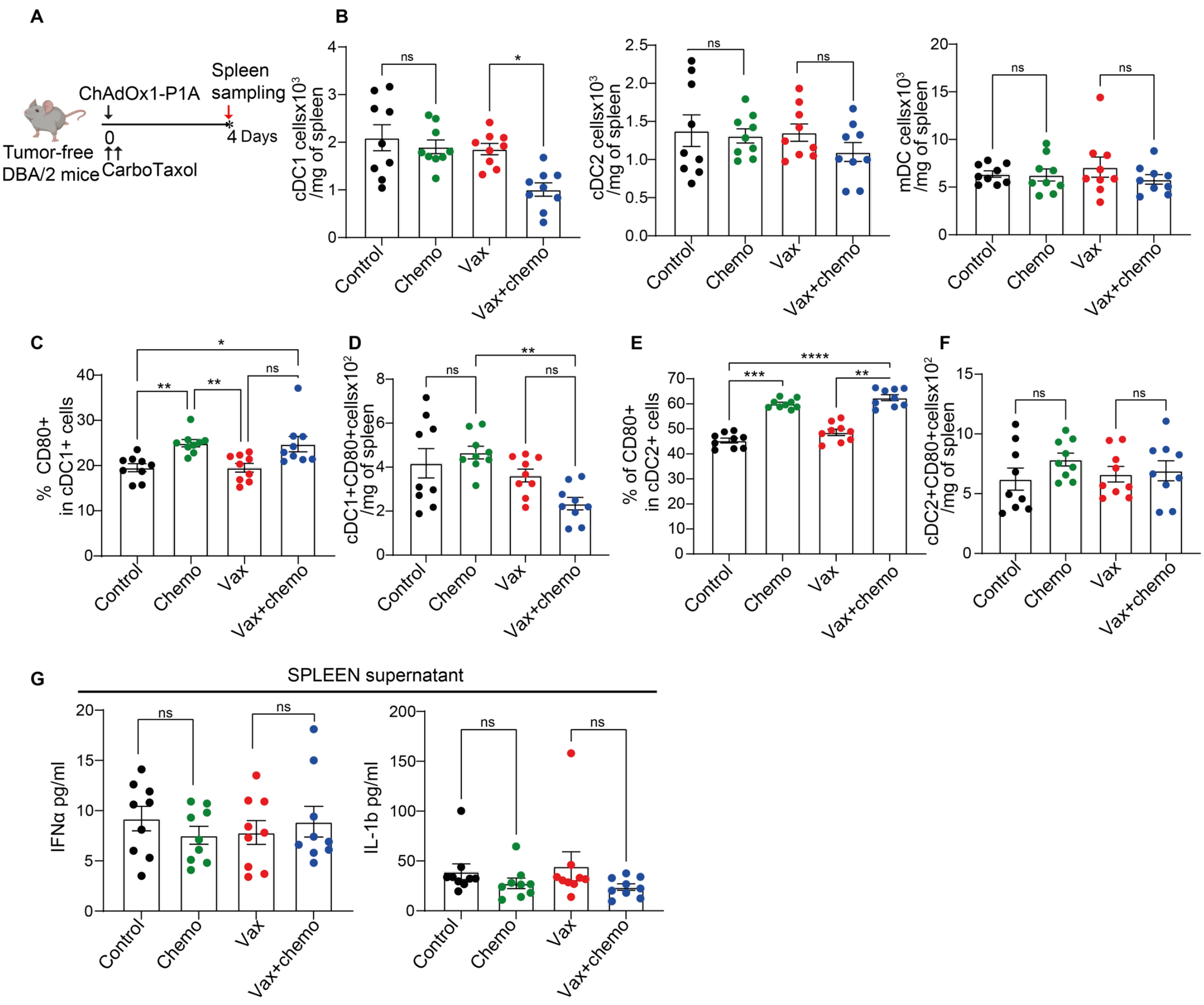


**Supplemental Figure 5: CarboTaxol does not impact DC subsets**

(**A**) Tumor free DBA/2 mice were vaccinated with ChAdOx1-P1A or PBS sham and treated with CarboTaxol. On day 4 post treatment, spleen was harvested as well as processing supernatant. Cells were stained and analysed by flow cytometry whilst supernatant were analysed by ELISA. Monocytic DC (mDC, defined as CD45+ Ly6G- F4/80+ CD11c+MHC2+), conventional DC1 (cDC1, defined as CD45+ Ly6G- F4/80- CD11c+ MHC2+ XCR1+) and conventional DC2 (cDC2, defined as CD45+ Ly6G- F4/80- CD11c+ MHC2+ CD172+) as well as the activation marker CD80 were evaluated.

(**B**) The cell numbers of cDC1, cDC2, mDC are shown.

(**C-D**) The proportion (**C**) and cell numbers (**D**) of activated cDC1 are shown.

(**E-F**) The proportion (**E**) and cell numbers (**F**) of activated cDC2 are shown.

(**G**) The level of IFN$\alpha$ and IL-1$\beta$ in the spleen are shown. Data are pooled from at least 2 independent experiments and mean ± SEM for each group are shown (n= 9 mice per group). Statistically significant differences between groups were determined by a Kruskal-Wallis test with Dunn's multiple comparisons test. ns non-significant, * $p \leq 0.05$, ** $p \leq 0.01$, *** $p \leq 0.001$, **** $p \leq 0.0001$

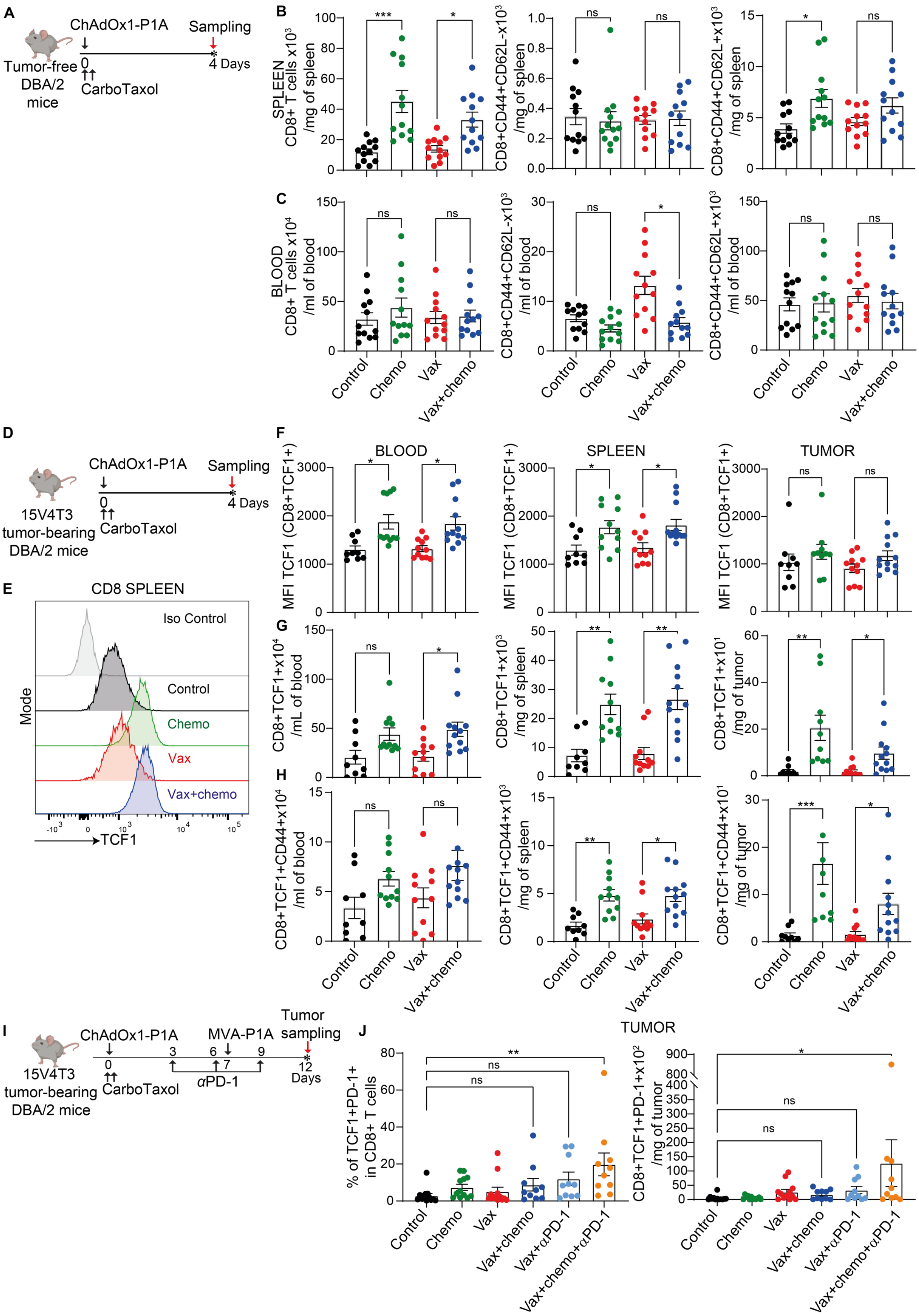
A
ChAdOx1-P1A
Sampling
Tumor-free DBA/2 mice
CarboTaxol
4 Days
B
SPLEEN CD8+ T cells x10^3 /mg of spleen
CD8+CD44+CD62L-x10^3 /mg of spleen
CD8+CD44+CD62L+x10^3 /mg of spleen
C
BLOOD CD8+ T cells x10^4 /ml of blood
CD8+CD44+CD62L-x10^3 /ml of blood
CD8+CD44+CD62L+x10^3 /ml of blood
Control
Chemo
Vax
Vax+chemo
D
ChAdOx1-P1A
Sampling
15V4T3 tumor-bearing DBA/2 mice
CarboTaxol
4 Days
E
CD8 SPLEEN
Iso Control
Control
Chemo
Vax
Vax+chemo
Mode
TCF1
F
BLOOD
SPLEEN
TUMOR
MFI TCF1 (CD8+TCF1+)
G
CD8+TCF1+x10^4 /mL of blood
CD8+TCF1+x10^3 /mg of spleen
CD8+TCF1+x10^1 /mg of tumor
H
CD8+TCF1+CD44+x10^4 /ml of blood
CD8+TCF1+CD44+x10^3 /mg of spleen
CD8+TCF1+CD44+x10^1 /mg of tumor
I
ChAdOx1-P1A
MVA-P1A
Tumor sampling
15V4T3 tumor-bearing DBA/2 mice
CarboTaxol
αPD-1
12 Days
J
TUMOR
% of TCF1+PD-1+ in CD8+ T cells
CD8+TCF1+PD-1+x10^2 /mg of tumor
Control
Chemo
Vax
Vax+chemo
Vax+αPD-1
Vax+chemo+αPD-1

**Supplemental Figure 6: CD8+ cell phenotype in tumor-free and tumor-bearing mice.**
(**A**) Scheme of treatment for tumour free DBA/2 mice.
(**B**) 4 days post treatment numbers of CD8, effector memory CD8 and central memory CD8 are shown in the spleen as mean ± SEM.
(**C**) Numbers of CD8, effector memory CD8 and central memory CD8 are shown in the blood as mean ± SEM. For (H-I) data are pooled from 3 independent experiments (n= 12 mice per group).
(**D**) Scheme of treatment for 15V4T3 tumor-bearing DBA/2. Mice were vaccinated with ChAdOx1-P1A or PBS sham and treated with CarboTaxol. At day 4 post treatment, spleen, blood and tumor were harvested.
(**E**) The level of TCF1 expression in CD8+ T cells in the spleen is shown.
(**F**) The MFI of TCF1+ T cells is shown in blood, spleen and tumor.
(**G**) The numbers of CD8+TCF1+ cells are shown in the blood, spleen and tumor.
(**H**) The numbers of CD8+CD44+TCF1+ cells are shown in the blood, spleen and tumor. The data are representative of 3 independent experiments and mean ± SEM for each group is shown (n=9 to 12 mice per group).
(**I**) Scheme of treatment for 15V4T3 tumor-bearing DBA/2 mice. To determine the TCF1 expression maintenance, tumor of each treatment group were harvested on day 12 post treatment.
(**J**) Proportion of TCF1+PD1+ in CD8+ T cells and cell numbers are shown as mean ± SEM. The data are pooled from 2 independent experiments (n=10 to 12 mice per group).
Statistically significant differences between groups were determined by a Kruskal-Wallis test with Dunn's multiple comparisons test. ns non-significant, * $p \leq 0.05$, ** $p \leq 0.01$, *** $p \leq 0.001$, **** $p \leq 0.0001$.

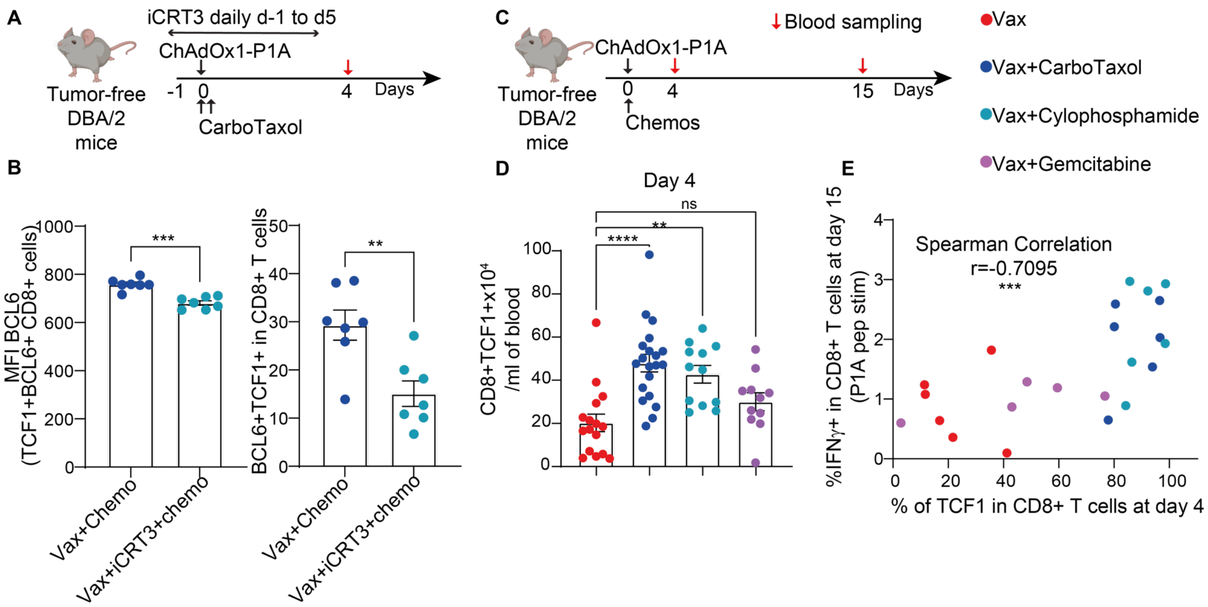


**Supplemental Figure 7: Importance of TCF1 expression for chemotherapy mediated-adjuvant effects**.

(**A**) Tumor free DBA/2 mice were vaccinated with ChAdOx1-P1A or PBS sham and treated with CarboTaxol on the same day. Mice were additionally treated with iCRT3 (20mg/kg, 40mg/kg on day 3) daily for 6 days starting one day prior to the prime. Blood was harvested and TCF1 expression as well as its transcriptional target BCL6 were investigated.

(**B**) The MFI level of BCL6 in TCF1+BCL6 in $CD8^{+}$ T cells and the proportion of TCF1+BCL6+ cells in CD8+T cells are shown. Statistically significant differences between groups were determined by a Mann-Whitney test.

(**C**) Tumor free DBA/2 mice were vaccinated with ChAdOx1-P1A or PBS sham and treated with CarboTaxol or cyclophosphamide or gemcitabine on the same day.

(**D**) Absolute numbers of CD8+TCF1+ T cells in the blood on day 4 post treatment are shown as mean ± SEM. Statistically significant differences between groups were determined by a Kruskal-Wallis test with Dunn's multiple comparisons test.

(**E**) TCF1 expression in CD8+ T cells as well as INFγ+P1A specific CD8+ T cells were evaluated at day 4 and 15 post ChAdOx1-P1A respectively. The correlation between these two populations is shown. Significance was evaluated using a spearman correlation test. ns non-significant, * $p \leq 0.05$, ** $p \leq 0.01$, *** $p \leq 0.001$, **** $p \leq 0.0001$
